# Inference of Nonlinear Spatial Subunits in Primate Retina with Spike-Triggered Clustering

**DOI:** 10.1101/496422

**Authors:** Nishal P. Shah, Nora Brackbill, Colleen E. Rhoades, Alexandra Kling, Georges Goetz, Alan Litke, Alexander Sher, Eero P. Simoncelli, E.J. Chichilnisky

## Abstract

Integration of rectified synaptic inputs is a widespread nonlinear motif in sensory neuroscience. We present a novel method for maximum likelihood estimation of nonlinear subunits by soft-clustering spike-triggered stimuli. Subunits estimated from parasol ganglion cells recorded in macaque retina partitioned the receptive field into compact regions, likely representing bipolar cell inputs. Joint clustering with multiple RGCs revealed shared subunits in neighboring cells, producing a parsimonious population model. Closed-loop subunit validation was then performed by projecting white noise into the null space of the linear receptive field. Responses to these null stimuli were more accurately explained by a model with multiple subunits, and were stronger in OFF cells than ON cells. Presentation of natural stimuli containing jittering edges and textures also revealed greater response prediction accuracy with the subunit model. Finally, the generality of the approach was demonstrated by application to V1 data.

## Introduction

The functional properties of sensory neurons are often probed by presenting stimuli and then inferring the rules by which inputs from presynaptic neurons are combined to produce receptive field selectivity. The most common simplifying assumption in these models is that the combination rule is *linear*, but it is well-known that many aspects of neural response are highly nonlinear. For example, retinal ganglion cells (RGCs) are known to be driven by nonlinear “subunits” (Hochstein and Shapley, 1976), which reflect signals from presynaptic bipolar cells that are rectified at the synapse onto RGCs (Demb et al., 1999, 2001; Borghuis et al., 2013). This subunit computation is fundamentally nonlinear because hyperpolarization of one bipolar cell input does not cancel depolarization of another. The subunit architecture endows RGCs with sensitivity to finer spatial detail than would arise from a linear receptive field (Hochstein and Shapley, 1976; Lee et al., 1995; Demb et al., 2001; Baccus et al., 2008; Crook et al., 2008; Schwartz et al., 2012), and has been implicated in the processing of visual features like object motion and looming (Olveczky et al., 2007; Munch et al., 2009). As another example, complex cells in primary visual cortex are thought to perform subunit computations on simple cell inputs, producing invariance to the spatial location of a stimulus. Indeed, subunits appear to be a common computational motif in the brain, perhaps because low maintained firing rates are common, and because the relation between depolarization and calcium entry that drives synaptic release is strongly rectifying. Thus, inferring neural circuitry from measurements of receptive field selectivity requires approaches that account for nonlinear computations such as those performed by subunits.

To this end, simple techniques have been used to reveal the presence and typical sizes of subunits (Hochstein & Shapley 1976), and more elaborate and/or ad hoc techniques have been used in specific experimental settings (Emerson et al., 1992; Paninski 2003; Sharpee et al., 2004; Rust et al., 2005; Schwartz et al., 2006; Pillow et al., 2006; Rajan & Bialek, 2013; Park et al, 2013; Theis et al, 2013; Liu et al., 2017; Freeman et al., 2015; Maheswaranathan et al., 2018; Shi et al., 2018). However, no widely accepted general computational tools exist for inferring the structure of nonlinear subunits providing inputs to a neuron, such as their individual sizes, shapes, and spatial arrangement. Because subunit computations are common, this lack of computational tools limits our ability to reason about neural circuitry in many parts of the nervous system.

Here, we develop a novel method for estimating the properties of subunits providing inputs to a neuron, and use it to probe subunit structure in parasol retinal ganglion cells (RGCs) in the primate retina. The method is derived as a maximum likelihood procedure for estimating subunit model parameters, and takes the form of a spike-triggered clustering algorithm. As such, it provides a natural generalization of spike-triggered averaging, which is widely used to estimate linear receptive fields because of their conceptual simplicity, robustness, and straightforward interpretation. In populations of recorded RGCs, the new method reveals a gradual partitioning of receptive fields with a hierarchical organization of spatially localized and regularly spaced subunits, consistent with the input from a mosaic of bipolar cells, and generalizes naturally and parsimoniously to the RGC population. A novel closed-loop stimulus reveals that RGC responses are explained substantially more accurately by the subunit model than by a linear model, a result that is stronger in OFF cells than ON cells, consistent with their known stronger synaptic nonlinearity, and that extends to naturalistic stimuli. Application of the method to primary visual cortex reveals subunits with expected spatiotemporal structure, suggesting that the approach will generalize to other contexts and neural circuits.

## Results

Our goal is to develop a general model for subunit computations in neural responses along with a methodology for parameter estimation, to test whether this model can predict responses over a wide range of stimuli, and to investigate the spatial properties of the estimated subunits in the model.

### Subunit response model and parameter estimation

In the model, RGC light responses are described as an alternating cascade of two linear-nonlinear (LN) stages (Figure 1A), a common functional form. In the first LN stage, the image intensities over space and recent time are weighted and summed by linear subunit filters, followed by an exponential nonlinearity. In the second LN stage, the subunit outputs are weighted by non-negative scalars, summed, and passed through a final output nonlinearity. This yields the neuron’s firing rate, which drives a Poisson spike generator.

**Figure 1:**
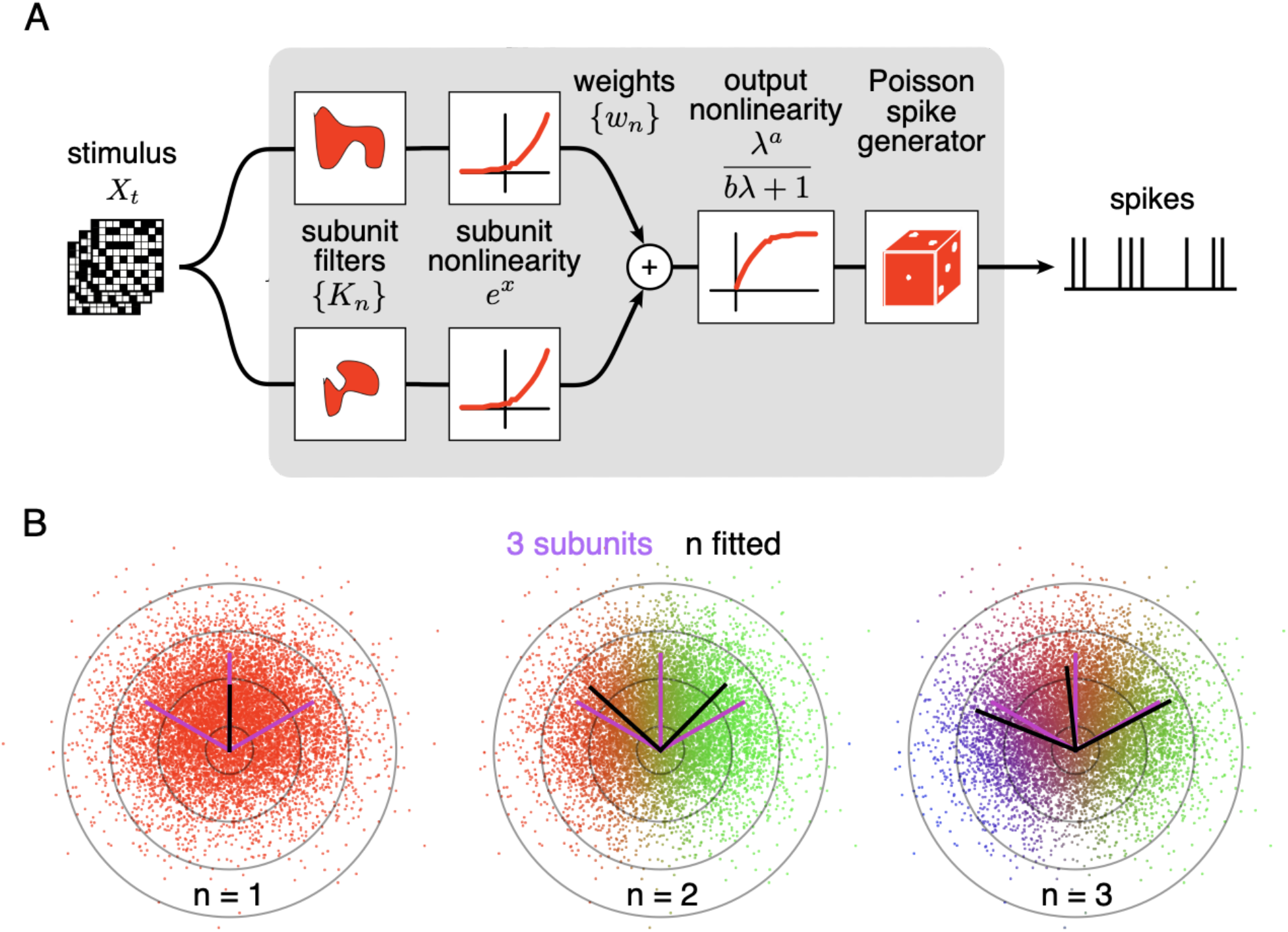
Spiking response model, and estimation through spike-triggered stimulus clustering. (A) The model is constructed as a cascade of two linear-nonlinear (LN) stages. In the first stage, subunit activations are computed by linearly filtering the stimulus (X_t_) with kernels (K_n_) followed by an exponential nonlinearity. In the second stage, a sum of subunit activations, weighted by (w_n_), is passed through a saturating output nonlinearity (g), yielding a firing rate that drives an inhomogeneous Poisson spike generator. (B) Estimation of subunits in simulated 3-subunit model cell by clustering of spike-triggered stimuli. Responses are generated to two-dimensional Gaussian stimuli (iso-probability circular contours shown) with three subunits (magenta lines), generating a spike-triggered stimulus ensemble (colored dots). Weights for different subunits are assigned to each spike triggered stimulus, with colors indicating the relative weights for different subunits. Subunits are then estimated by weighted summation of spike triggered stimulus ensemble (black lines). Soft-clustering of spike triggered stimuli with different number of clusters results in progressive estimation of the underlying subunits. Clustering with correct number of subunits (right panel) results in estimated subunits (black lines) aligned with the true subunits.

To predict neural responses using the model, the model parameters (subunits, weights, and nonlinearity) must be estimated from recorded data. However, the estimation problem is difficult because the negative log-likelihood is not a convex function of the parameters. We derive an efficient solution that operates by alternating between two steps (see Methods): (1) estimate the subunits by identifying clusters in the space of spike-triggered stimuli, in a way that follows naturally from maximizing likelihood (Figure 1B), and (2) estimate the parameters of the output nonlinearity using standard optimization methods. This *spike-triggered clustering* solution offers a simple and robust generalization of the commonly-used spike-triggered averaging method for receptive field estimation. Intuitively, clustering is effective because stimuli that produce spikes will typically fall into distinct categories based on which subunit(s) they activate.

Simulations with a realistic RGC model show that this procedure yields accurate estimates of model parameters when sufficient data are available (Figure S1). In addition, when experimental data are limited, reliable subunit estimates are attained by regularizing the likelihood objective to capture prior knowledge about the spatial localization of subunits, enforced as a projection step in the clustering loop (see Methods, Figure 3). Note that the model includes two hyperparameters (number of subunits and regularization strength) that are set by optimizing model predictions on held-out data (i.e., cross-validation).

### Estimated subunits are spatially localized and non-overlapping

To test the subunit model and estimation procedure on primate RGCs, light responses were obtained using large-scale multielectrode recordings from isolated macaque retina (Litke et al., 2004, Frechette et al., 2005). Spiking responses of hundreds of RGCs to a spatiotemporal white noise stimulus were used to classify distinct cell types, by examining properties of the spike-triggered average stimulus (STA), which provides a linear summary of light response (Frechette et al, 2005; Chichilnisky et al., 2002; Field et al; 2007). Complete populations of ON and OFF parasol RGCs covering the recorded region were examined further.

Fitting a subunit model reliably to these data was aided by decoupling the spatial and temporal properties of subunits. Specifically, although the fitting approach in principle permits estimation of full spatiotemporal subunit filters, the high dimensionality of these filters requires substantial data (and thus, long recordings). The parameter space was reduced by assuming that subunit responses are spatio-temporally separable, and that all subunits have the same temporal filter (consistent with known properties of retinal bipolar cells that are thought to form the RGC subunits). Given these assumptions, the temporal filter was estimated from the STA (see Methods). The stimulus was then convolved with this time course to produce an instantaneous spatial stimulus associated with each spike time. This temporally prefiltered spatial stimulus was then used to fit a model with purely spatial subunits.

The number of subunits, N, is a discrete parameter, and we examined its effect by fitting the model for different N, each with an independent initialization. Setting N=1 yielded a single subunit whose receptive field corresponds to a LN model. Typically, a model with two subunits partitioned this receptive field into spatially localized regions (Figure 2A, second row). Fitting the model with 3 or more subunits (Figure 2A, subsequent rows) typically caused one of the subunits from the preceding row to be partitioned further, while other subunits were largely unchanged. In principle this procedure could be repeated many times to capture all subunits in the receptive field. However, the number of model parameters increases with N, and the estimation accuracy decreases, with estimated subunits becoming noisier with larger overlap (Figure 2A, last row). This observation suggests an estimation approach in which the hierarchical introduction of subunits is built into the procedure, with the potential for greater efficiency (see Methods and Figure S2).

**Figure 2:**
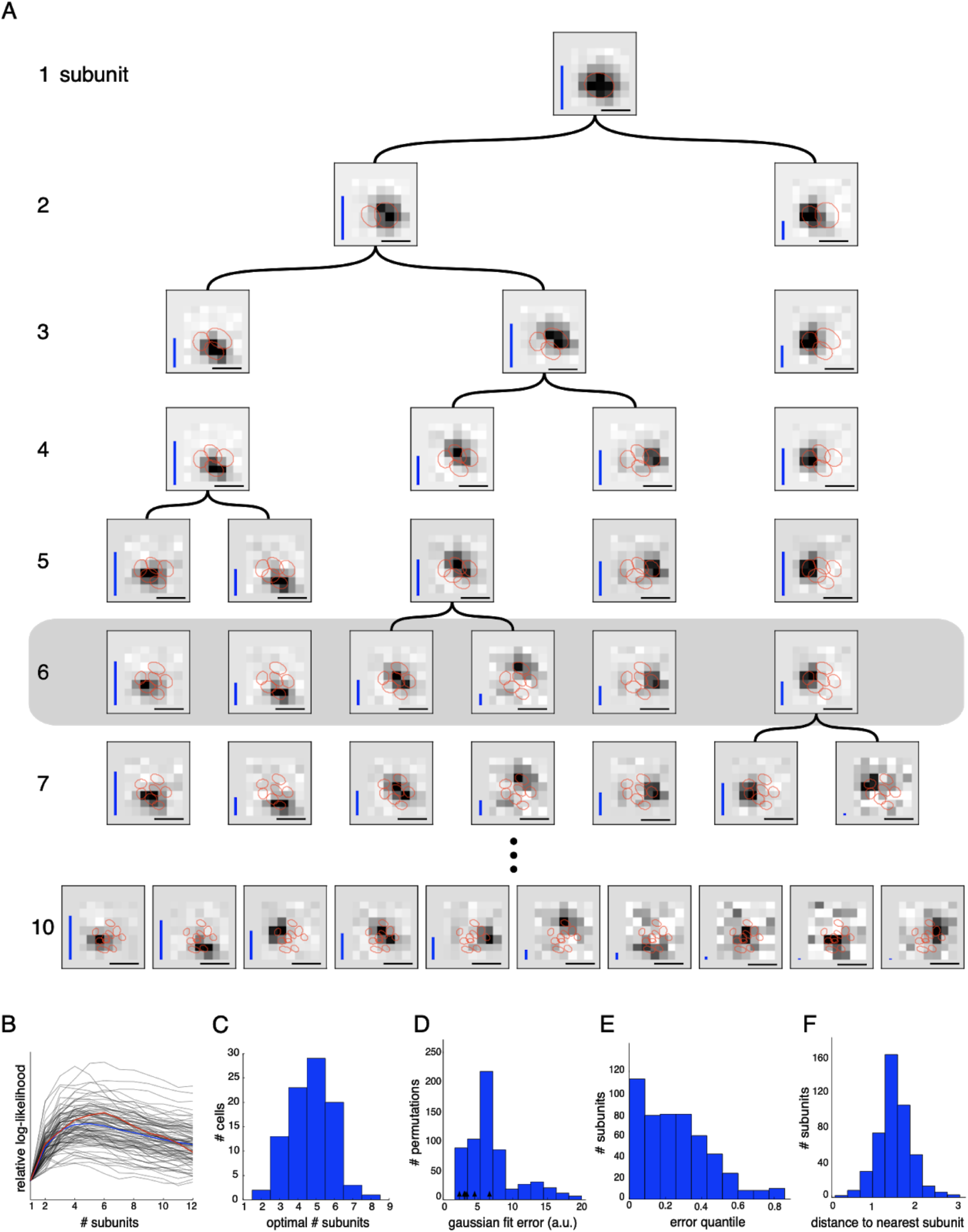
Estimated subunit properties. (A) Subunits, shown as grayscale images, estimated from OFF parasol cell responses to 24 min of white noise. Each pixel was temporally prefiltered with a kernel derived from the spike-triggered average (STA). Rows show estimated spatial subunits for successive values of N. Subunit locations are indicated with red ellipses, corresponding to the contour of a fitted two-dimensional Gaussian with standard deviation equal to the average nearest neighbor separation between subunits. As N increases, each successive set of subunits may be (approximately) described as resulting from splitting one subunit into two (indicated by arrows). Large N (e.g., last row) yields some subunits that are noisy or overlap substantially with each other. Height of vertical blue bars indicate the relative strength (average contribution to response over stimulus ensemble) of each subunit. Horizontal black bars indicate spatial scale (150μm). (B) Log-likelihood as a function of number of subunits (relative to single subunit model) for 91 OFF parasol cells (black) on 3 min of held-out test data, averaged across 10 random initializations of the model, from a distinct randomly sampled training data (24 min from remaining 27 min of data). Population average is shown in blue and the example cell from (A) is shown in red. (C) Distribution of optimal number of subunits across different cells, as determined by cross-validated log-likelihood on a held-out test set for OFF parasol cells. (D, E) Spatial locality of OFF parasol subunits, as measured by mean-squared error of 2D gaussian fits to subunits after normalizing with the maximum weight over space. Control subunits are generated by randomly permuting pixel weights for different subunits within a cell. For this analysis, the optimal number of subunits was chosen for each cell. (D) Distribution of MSE values for randomly permuted subunits for the cell shown in (A). MSE of 6 (optimal N) estimated subunits indicated with black arrows. (E) Distribution of quantiles of estimated OFF parasol subunits, relative to the distribution of MSE values for permuted subunits, across all cells and subunits. Null hypothesis has uniform distribution between 0-1. (F) Distribution of distances to nearest neighboring subunit within each OFF parasol cell. Distances are normalized by geometric mean of standard deviation of the gaussian fits along the line joining center of subunits. For this analysis, each cell is fit with 5 subunits (most frequent optimum from (C)).

An optimal number of subunits was chosen for each cell as the *N* that maximized the cross-validated likelihood (i.e., the likelihood measured on a held out test set - Figure 2B). The typical optimum for OFF cells was 4-6 subunits (Figure 2C), a value that is governed by the true number of subunits, the resolution of the stimulus (which governs the dimensionality of the parameter space, see Figure 3), and the amount of data (number of spikes). Since the ON parasol cells had smaller optimum number of subunits (1-2) and much smaller increases in likelihood of the subunit model compared to an LN model (not shown), we focus subsequent analysis only on the OFF parasol cells.

**Figure 3:**
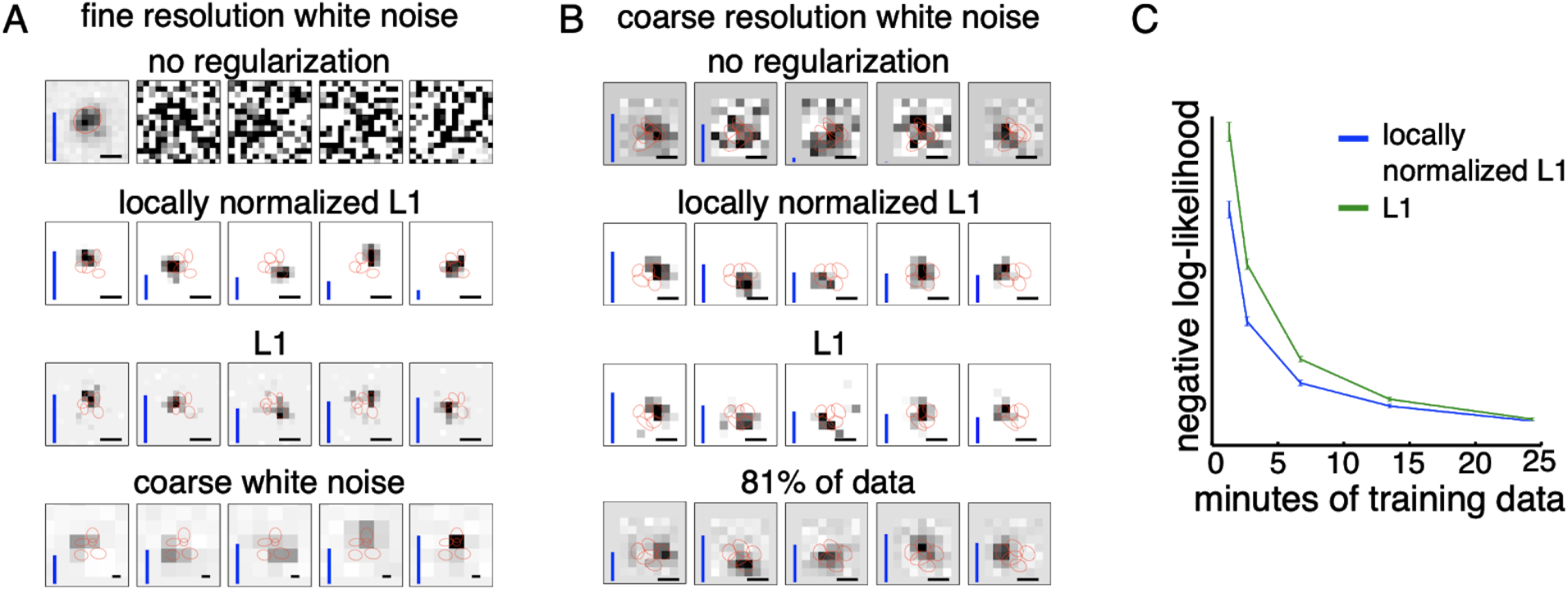
Spatially localized subunit estimation. Comparison of different priors for estimating subunits using limited data. Examples of OFF parasol cells are shown. (A) Five subunits estimated using all data (1 hour 37 min) for fine resolution white noise without regularization (top row). The first estimated subunit is identical to the full receptive field and the others are dominated by noise. Locally normalized L1 (second row) and L1 (third row) regularization both give spatially localized subunits, with L1 regularization leaving more noisy pixels in background. In both cases, optimal regularization strength was chosen (from 0-1.8, steps of 0.1) based on performance on held-out validation data. The contours reveal interdigitation of subunits (red lines). Subunits estimated using white noise with 2.5x coarser resolution (24 min) and no regularization are larger, but at similar locations as subunits with fine resolution (bottom row). Scale bar: 75μm (B) For the cell in Figure 1A, 5 subunits estimated using the 3 min (10% of recorded data) of coarse resolution white noise responses are noisy and non-localized (top row). Similar to the fine case, using locally normalized L1 (second row) and L1 (third row) regularization both give spatially localized subunits, with L1 regularization subunits having noisier background pixels. The regularization strength (between 0 - 2.1, steps of 0.1) was chosen to maximize log-likelihood on held out data (last 5 min of data). Subunits estimated using 24 min (81% of data) of data are spatially localized and partition the receptive field (bottom row). Scale bars: 150μm (C) Held out log-likelihood for a 5 subunit model estimated from varying durations of training data with L1 (green) and locally normalized L1 (blue) regularization. Results averaged over 91 OFF parasol cells from Figure 1A.

The estimated subunits were larger and fewer in number than expected from the size and number of bipolar cells providing input to parasol RGCs at the eccentricities recorded (Jacoby et al., 2000; Schwartz & Rieke 2011; Tsukamoto & Omi 2015). However, two other features of estimated subunits suggested a relationship to the collection of bipolar cells contributing to the receptive field. First, estimated subunits were compact in space as shown in Figure 2A. This was quantified by comparing each subunit with the collection of subunits derived by randomly permuting the filter values at each pixel location across different subunits within a cell. Spatial locality for each subunit was evaluated by the mean-squared error (MSE) of 2D Gaussian fits. Compared to the permuted subunits, the estimated subunits had substantially lower error (Figure 2D, E). Second, the subunits “tiled” the RGC receptive field, in that they covered it with only small amounts of overlap, while leaving no holes. This was quantified by noting that on average, the neighboring subunits for a given cell were separated by ~1.5 times their width, with relatively little variation (Figure 2F).

Given these observations, a natural hypothesis for subunits observed with coarse stimulation is that they correspond to aggregates of neighboring bipolar cells. To test this intuition, the algorithm was applied to white noise responses generated from a simulated RGC with a realistic size and number of subunits in the receptive field center (Jacoby et al., 2000; Schwartz & Rieke 2011). For data simulated using coarse pixelated stimuli matched to those used in the experiments, the estimated subunits were fewer and larger than the bipolar cells, with each subunit representing an aggregate of nearby underlying bipolar cells (Figure S1). The recoveredd subunits also exhibited spatial locality and tiling. Thus, the simulation supports the hypothesis that the subunits estimated from recorded data (Figure 2A) reflect aggregates of adjacent underlying bipolar cells.

### Regularization for spatially localized subunit estimation

To estimate subunits with a resolution approaching that of the underlying biology would require higher resolution stimuli. But higher resolution stimuli typically require longer duration of data. Specifically, to obtain a constant quality estimate of subunits as the stimulus resolution/dimensionality is increased requires a proportional increase in the number of spikes (e.g. see Dayan & Abbott, 2001). Furthermore, higher resolution stimuli typically produce lower firing rates, because the effective stimulus contrast driving the photoreceptors is reduced. To illustrate the problem, the impact of increased resolution and smaller duration of data were examined separately. First, higher resolution stimuli led to poor subunit estimates, even when all the recorded data (97 minutes) were used (Figure 3A, top row): of the five estimated subunits, one resembled the receptive field, and the others were apparently dominated by noise and had low weights; thus, the algorithm effectively estimated a LN model. Second, for coarse resolution stimuli, using only 10% of the recorded data led to noisy subunit estimates (Figure 3B, top row) compared to the estimates obtained with 81% of data (24 minutes, Figure 3B, last row). These observations suggest that it would be useful to incorporate prior knowledge about subunit structure, and thus potentially obtain more accurate estimation with limited data.

To accomplish this, a prior on subunit filters was introduced, based on the observed spatial locality of estimated subunits (Figure 2). Previous studies (Liu et al., 2017; Maheswaranathan et al., 2018; MacFarland et al., 2013), have used L1 regularization (a common means of inducing sparsity) but this is agnostic to spatial locality (and indeed is invariant to spatial rearrangement). Instead of L1 regularization, we developed a novel locally normalized L1 (LNL1) regularizer that penalizes large weights, but only if all of the neighboring weights are small: 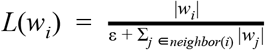 for ε = 0.01. This penalty is designed to have a relatively modest influence on subunit size (compared to L1, which induces a strong preference for smaller subunits). The LNL1 penalty was incorporated using the projection operator in the clustering loop (Figure 1B). In this case, the appropriate projection operator is an extension of the soft-thresholding operator for the L1 regularizer, where the threshold for each pixel depends on the strength for neighboring pixels. The regularization strength is chosen to maximize cross-validated likelihood (see Methods).

The proposed prior improved subunit estimates for both of the limited data settings (high resolution, or small duration) presented above. In the case of the coarse resolution data, both L1 and LNL1 priors produced spatially localized subunits (Figure 3B, middle rows) whose locations matched the location of the subunits estimated using more data without regularization (Figure 3B, last row). This suggests that the proposed prior leads to efficient estimates without introducing new biases. Relative to L1 regularization, LNL1 regularization yielded fewer apparent noise pixels in the background (Figure 3B, middle rows). This improvement in spatial locality is reflected in more accurate response prediction with LNL1 regularization (Figure 3C).

In the case of high dimensional stimuli (Figure 3A, top row), both L1 and LNL1 priors were successful in estimating compact subunits (Figure 3A, middle rows) and matched the location of subunits obtained with a coarser stimulus (Figure 3A, bottom row). The estimated subunits tiled the receptive field, as would be expected from an underlying mosaic of bipolar cells. The LNL1 prior also led to subunit estimates with fewer spurious pixels compared to the L1 prior. Note that although these spurious pixels can also be suppressed by using a larger weight on the L1 prior, this approach led to smaller subunit size, which produced less accurate response predictions, while LNL1 showed a much smaller dependence of subunit size (not shown). Hence, in both the conditions of limited data examined, the novel LNL1 prior yielded spatially localized subunits, fewer spurious pixels, and a net improvement in response prediction accuracy.

### Parsimonious modeling of population responses using shared subunits

To fully understand the visual processing in retina it is necessary to understand how a population of RGCs coordinate to encode the visual stimulus in their joint activity. The subunit estimation method extends naturally to joint identification of subunits in populations of neurons. Since neighboring OFF parasol cells have partially overlapping receptive fields (Gauthier et al., 2009), and sample a mosaic of bipolar cells, neighboring cells would be expected to receive input from common subunits. Indeed, in recorded OFF parasol mosaics, estimated subunits were frequently closer together than expected from the distribution of nearest-neighbor subunit distances obtained from single-cell data. For example, in one data set, the fraction of nearby subunits from different cells (spaced by less than 1SD of a Gaussian fit) was 14%, substantially higher than the predicted value of 9% based on the observed separation of subunits within a cell (Figure 2F). Examination of the mosaic suggests that these pairs of closely-spaced subunits may in fact correspond to single subunits shared between neighboring RGCs (Figure 4A). This raises the possibility that the estimated subunits of neighboring cells often reflect the same underlying biological substrate, and that joint estimation of subunits from the pooled data of neighboring RGCs may more efficiently reveal their collective spatial structure.

**Figure 4:**
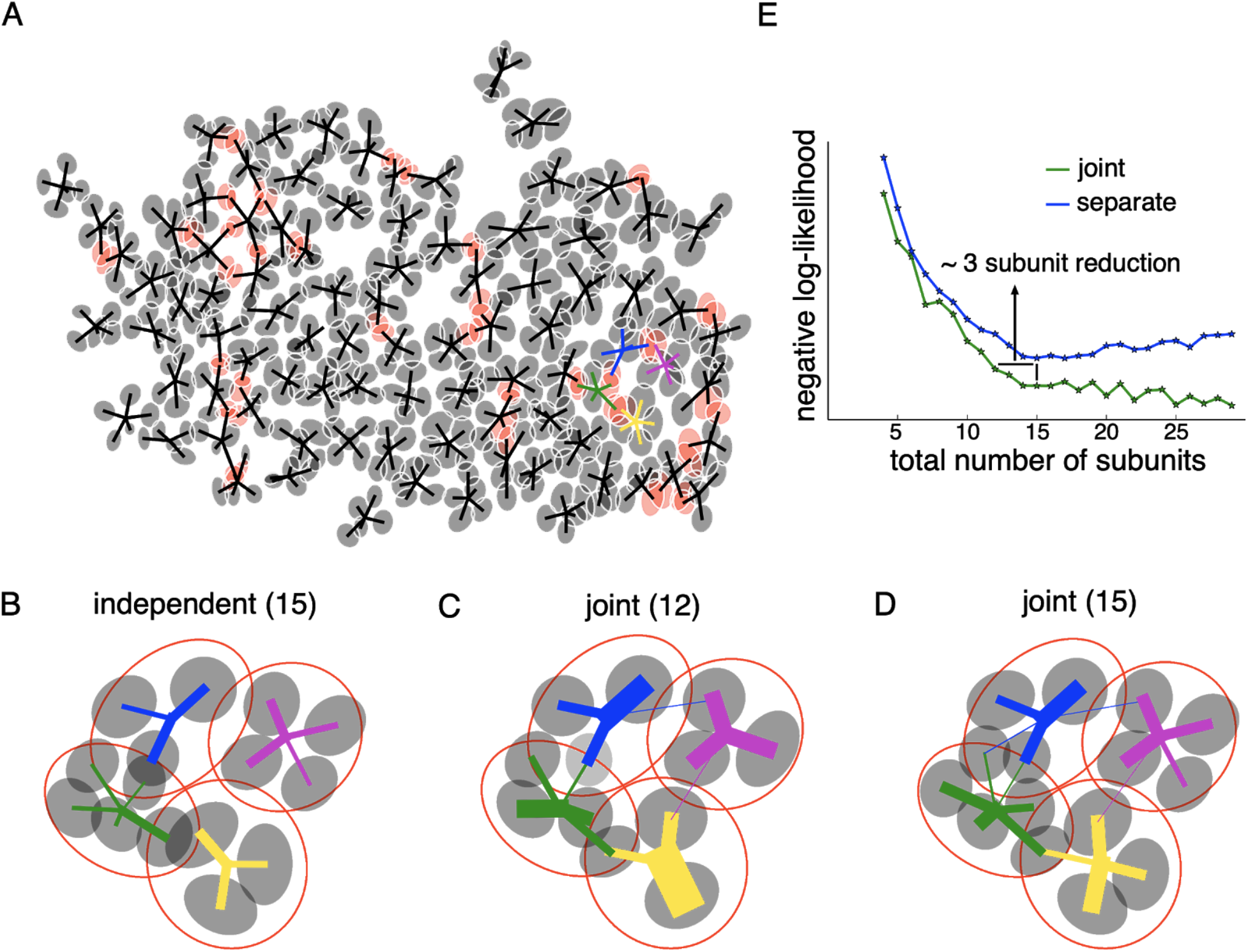
Joint estimation of subunits across multiple nearby cells. (A) Gaussian fits to the subunits estimated for an entire OFF parasol cell population (five subunits per cell, with poorly estimated subunits removed). Lines connect center of each cell to its subunits. Subunits which are closer to its neighbor than average (below 15 percentile) are indicated in red. (B) Different number of subunits for each cell is chosen (total 15 subunits) to give the highest summation of log-likelihood for four nearby cells. Gaussian fits to receptive field of the cells (red) and their subunits (gray). Connection strength from a cell (distinct color) to its subunits is indicated by thickness of line. (C) A common bank of 12 subunits estimated by jointly fitting responses for all four cells gives similar accuracy as (B). Sharing is indicated by subunits connected with lines of different colors (different cells) (D) A model with a common bank of 15 subunits (same number as B) gives better performance than than estimating the subunits for each cell separately. (E) The total negative log-likelihood on test data for four cells (y-axis) v/s total number of subunits (x-axis) for the two population models with subunits estimated jointly across cells (green) and best combination of separately estimated subunits (blue). Horizontal shift between curves indicates the reduction in total number of subunits by jointly estimating subunits for nearby cells for similar prediction accuracy. The vertical shift indicates better performance by sharing subunits, for a fixed total number of subunits. In each case, the best value for locally normalized L1 regularization (in 0-3, step size 0.2) for a fixed number of subunits was chosen based on the performance on a separate validation dataset (see Methods).

This hypothesis was tested by estimating a common “bank” of subunits to simultaneously predict the responses for multiple nearby cells. Specifically, the firing rate for the i^th^ neuron was modeled as: *R_i_* = *g_i_*(Σ*w_n,j_exp*(*K_n_* • *X_t_*)). Here X_t_ is the stimulus, K_n_ are the filters for the common population of subunits, g_i_ is the output nonlinearity for the i^th^ neuron,and w_n,i_ is the non-negative weight of subunit n to the i^th^ neuron, which is 0 if they are not connected. The model parameters were estimated by maximizing the summation of log-likelihood across cells, again in two steps. For the first step, the output nonlinearities were ignored for each cell and a common set of subunit filters were found, that clustered the spike-triggered stimuli for all the cells simultaneously. Specifically on each iteration, a) the relative weights for different subunits were computed for each spike-triggered stimulus and b) the cluster centers were updated using the weighted sum of spike-triggered stimuli across different cells (see Methods). Since the subunit filters could span the receptive fields of all the cells, leading to higher dimensionality, the spatial locality (LNL1) prior was used to efficiently estimate the subunits (Figure 3). In the second step of the algorithm, the non-linearities were then estimated for each cell independently (similar to Figure 1).

To quantify the advantages of joint subunit estimation, the method was applied to a group of 4 nearby OFF parasol cells. First, models with different number of subunits were estimated for each cell separately, and the combination of 15 subunits that maximized the cross-validated log-likelihood when summed over cells was chosen (Figure 4B). However, similar test log-likelihood was observed by using a joint model with only 12 subunits, effectively reducing model complexity by three subunits (Figure 4C). Examination of the subunit locations revealed that pairs of overlapping subunits from neighboring cells in separate fits were replaced with one shared subunit (Figure 4B, C). Also, a joint model with 15 subunits showed higher prediction accuracy than a model in which 15 subunits were estimated separately (Figure 4D).

This trend was studied for different number of subunits in the filter bank, by comparing the total negative log-likelihood for these four cells associated with the joint model (Figure 4E, green) and a model in which subunits were estimated for each cell independently (Figure 4E, blue). Joint fitting provided more accurate predictions than separate fits, when using a large (>12) total number of subunits. For a smaller number of subunits, a horizontal shift in the observed likelihood curves revealed the reduction in the number of subunits that was obtained with joint estimation, without a reduction in prediction accuracy. Overall, jointly estimating a common collection of subunits for nearby cells resulted in a more parsimonious explanation of population responses.

### Subunit model explains spatial nonlinearities revealed by null stimulus

To more directly expose the advantage of a subunit model over commonly used models of RGC light response, experiments were performed using a visual stimulus that elicits no response in a LN model. Contrast-reversing spatial grating stimuli centered on the RF is an example of one such visual stimulus (Hochstein & Shapley 1976; Demb et al., 1999). However, to avoid recruiting additional nonlinearities, we wanted to use a stimulus with spatio-temporal statistics very similar to the diverse (high-entropy) white noise stimuli used for model characterization. Therefore, a stimulus was obtained by manipulating white noise so as to cancel the contribution of the linear RF.

Specifically, the computation performed by a hypothetical LN neuron can be represented in a stimulus space in which each axis represents stimulus contrast at a single pixel location (Figure 5A). In this representation, the LN response is determined by the projection of the stimulus vector onto the direction of the neuron’s spatial RF. Thus, a “null stimulus” orthogonal to the RF can be computed by subtracting this projection from the original stimulus (Figure 5B). This null stimulus should yield zero response in a LN neuron. Note that although this approach is described above purely in the spatial domain, it generalizes to the full spatiotemporal RF, and the procedure based on the spatial RF alone will produce a null stimulus for a spatio-temporal linear filter that is space-time separable.

**Figure 5:**
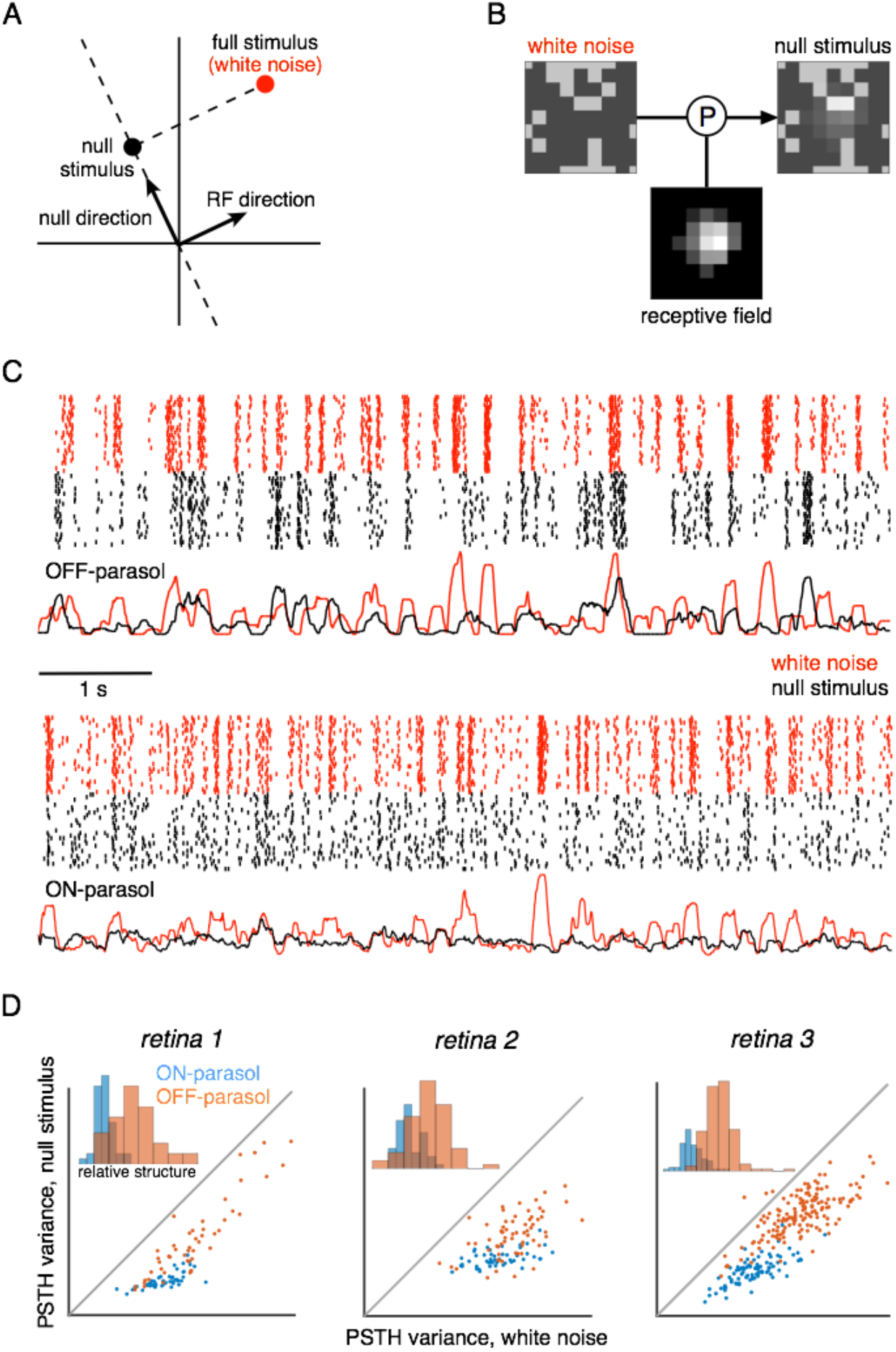
Cells respond to stimulus in null space of receptive field. (A) Construction of null stimulus, depicted in a 2-dimensional stimulus space. Each dimension of stimulus space consists of intensity along a particular pixel. A stimulus frame is represented as a point in this space. Each stimulus frame can be represented geometrically as the sum of the component along the receptive field (RF) direction (the response component of a linear nonlinear model) and the component orthogonal to the RF direction (the null component). (B) The null stimulus is constructed by projecting out the RF contribution from each frame of white noise stimulus. (C) Rasters representing responses for an OFF parasol (top) and ON parasol (bottom) cell to 30 repeats of 10s long white noise (red) and the corresponding null stimulus (black). (D) Response structure for ON parasol (blue) and OFF parasol (orange) populations for white noise (x-axis) and null stimulus (y-axis) across three preparations (different plots). The response structure was measured as variance of PSTH over time. Insets: Histogram of relative structure in white noise responses that is preserved in null stimulus for ON parasol (blue) and OFF parasol (orange). Relative structure is measured by ratio of response structure in the null stimulus and response structure in white noise stimulus.

We tested whether this null stimulus could silence the response of RGCs, as follows. A 10-sec movie of white noise stimulus frames was projected into the intersection of the spatial null spaces corresponding to all cells of a single type. Responses were recorded from 30 repeated presentations of both the white noise and the null stimulus. RGC firing rates showed a modulation for both white noise and the null stimulus that was highly reproducible across repeats (Figure 6C, D), indicating that the visual inputs are not integrated linearly across the RF. Note that the null stimulus modulated OFF parasol cells more strongly than ON parasol cells, consistent with previous results obtained with white noise and natural scene stimuli (Chichilnisky & Kalmar 2002, Demb et al., 2001, Turner & Rieke 2016); henceforth, analysis is focused on OFF parasol cells.

**Figure 6:**
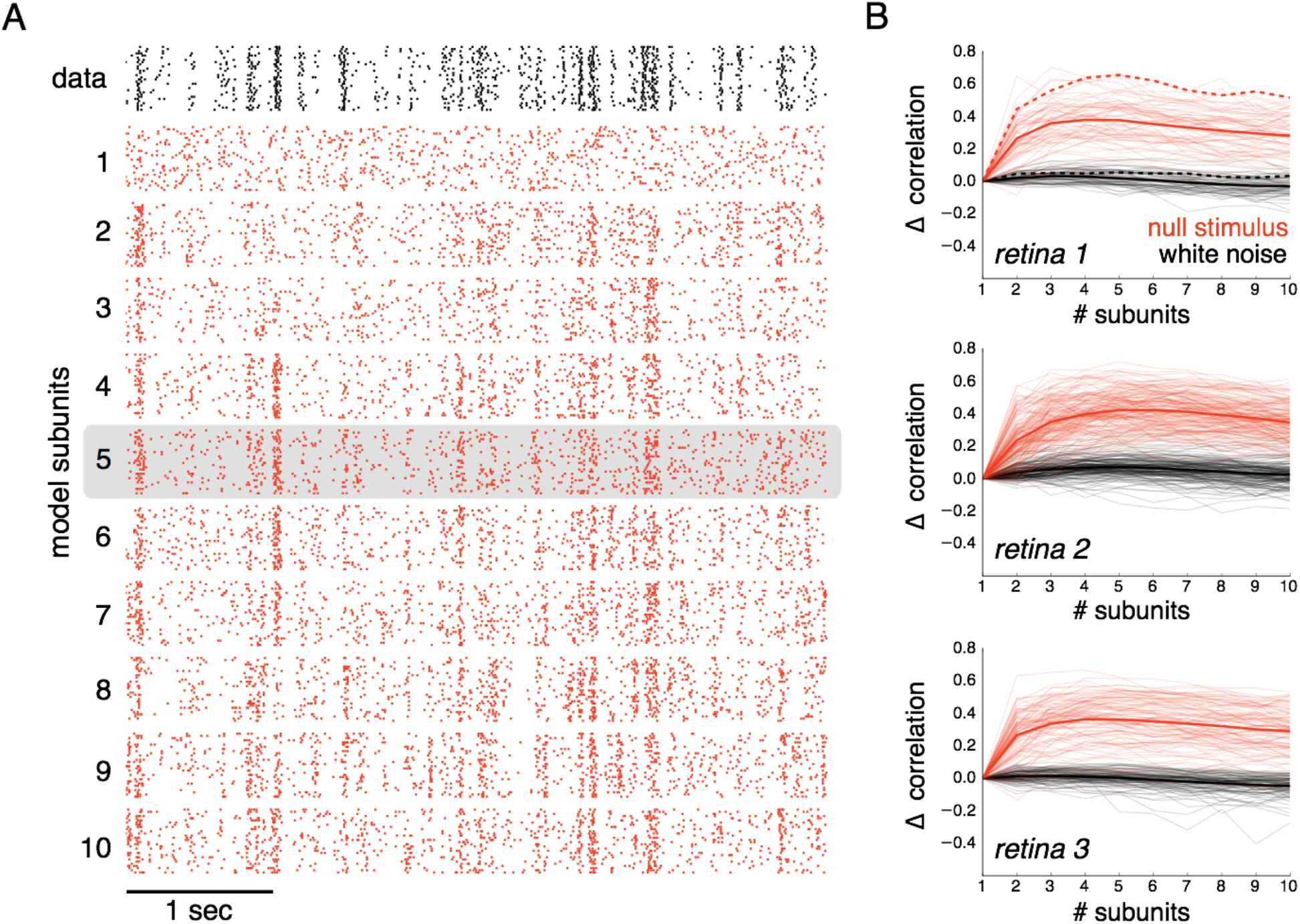
Subunits improve prediction of responses to null stimuli. (A) Rasters for recorded responses of an OFF parasol cell to 30 presentations of a 5 sec long null stimulus (top row, black). Predictions of models with increasing (1 to 10) number of subunits (subsequent rows, red). (B) The change in correlation between PSTH for recorded and predicted responses with different numbers of subunits across three preparations. Spatial kernels were estimated from 24 min of non-repeated white noise stimulus, with scales and output nonlinearity estimated from the first 5 sec of the repeated stimulus. Performance on the last 5 sec of the repeated stimulus was averaged over 10 fits, each with a random subsample of the non-repeated white noise stimulus. Individual OFF parasol cells (thin lines) and population average (thick lines) for the null stimulus (red) and white noise (black). The cell is (A) is indicated with dotted lines.

The subunit model captured a substantial component of response nonlinearities revealed by the null stimulus. Subunit filters were estimated from responses to white noise (step 1 of model estimation, Figure 1), and response nonlinearities were estimated from responses to null stimulus (step 2 of model estimation, Figure 1). As expected, the single subunit (LN) model failed to capture responses to the held-out null stimulus (Figure 6A, second row). As the number of subunits was increased, the model’s performance progressively increased (Figure 6A, last two rows). Prediction accuracy, defined as the correlation between the recorded and predicted firing rate, was evaluated across entire populations of OFF parasol cells in three recordings (Figure 6B). The single subunit model failed, with prediction accuracy near 0 (mean accuracy : 0.02, 0.12, 0.03 for three retinas). The prediction accuracy gradually increased with addition of more subunits, saturating at roughly 4-6 subunits (Figure 6B, red). In contrast, the addition of subunits showed marginal improvements for white noise stimuli, with signs of overfitting for large numbers of subunits (Figure 6B, black). Even though the log-likelihood increased for longer durations of a white noise stimulus (Figure 2B), the increase in correlation for repeats of shorter duration white noise was not significant. Thus, the response nonlinearities captured by the subunit model are more clearly exposed using the null stimulus.

### Subunits enhance response prediction for naturalistic stimuli

A fuller understanding of the biological role of spatial nonlinearities requires examining how they influence responses elicited by natural stimuli. Images from the Van Hateren database were shown for one second each, jittered according to the eye movement trajectories recorded during fixation by awake macaque monkeys (Z.M. Hafed and R.J. Krauzlis, personal communication; Heitmann et al, 2016, Van Hateren & van der Schaaf, 1998). The visual stimuli consisted of training data interleaved with identical repeats of testing data (see Methods). Subunit filters and time courses were first estimated using white noise stimuli (1st step, Figure 1), then the non-linearities and subunit magnitudes were fitted using natural stimuli (2nd step, Figure 1) (see Methods). Subsequently, the ability of the subunit model to capture responses to natural stimuli was measured.

Responses to natural stimuli showed a distinct structure, with strong periods of firing or silence immediately following saccades and occasional responses elicited by inter-saccadic eye-movements (Heitmann et al, 2016, Figure 7A black raster). This structure was progressively better captured by the model as the number of subunits increased, as shown by the rasters for an OFF parasol cell in Figure 7A. This finding was replicated in OFF parasol populations in three retinas, and quantified by the correlation between the recorded and predicted PSTH (Figure 7B). However, some of the firing events could not be captured even with a large number of subunits, suggesting additional mechanisms affecting natural scene responses.

**Figure 7:**
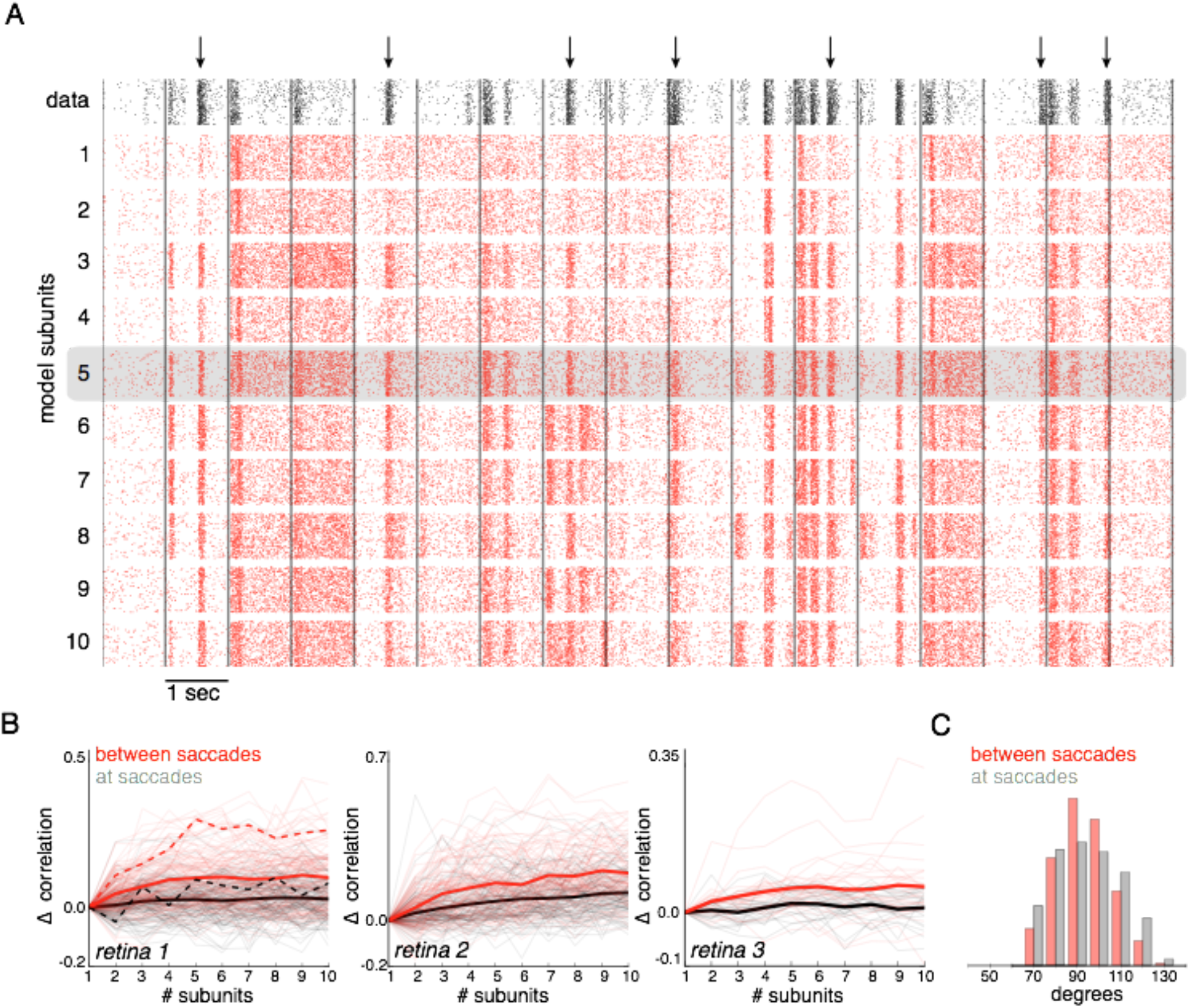
Subunits improve response prediction accuracy for NSEM. (A) Top row: Rasters of responses for an OFF parasol cell from 40 presentations for 30s long natural stimuli. Saccades to a new image every 1 sec (black lines). Subsequent rows indicate model predictions using different number of subunits (1 to 10) respectively. (B) Change in correlation between PSTH for recorded and predicted responses with different number of subunits across 3 preparations. Individual OFF parasol cells (thin lines) and population average (thick lines) for two conditions: 250 ms immediately following saccades (black) and inter-saccadic stimuli (red). Cell from (A) shown with dotted lines. (C) Distribution of angle between vectors representing the spatial receptive field and natural stimuli at saccades (black) and between saccades (red). Inter-saccadic stimulus shows more prevalence around 90 degrees (blue line).

Interestingly, further examination revealed that subunits led to greater improvement for responses elicited by inter-saccadic stimuli (small eye movements), compared to saccades (large eye movements) (Figure 7B). This could be explained by the fact that changes in luminance are more spatially uniform at saccades, resulting in more linear signaling by OFF parasol cells. Indeed, there were fewer stimuli in the null space of the receptive field (90 degrees) during saccades compared to between saccades (Figure 7C) and, the subunit model provides a more accurate explanation of responses to stimuli in the null space (Figure 6). Thus, these results indicate that spatial non-linearities explain the responses to natural stimuli, especially those elicited by jittering natural images.

### Application to simple and complex cells in primary visual cortex

Identification of subunits using spike-triggered stimulus clustering should generalize to other neural systems with similar cascade of linear and non linear operations. We applied the unregularized method to data obtained from V1 simple and complex cells, responding to spatio-temporally oriented flickering bars (Rust et al, 2005). The number of cross validated subunits for the complex cell was greater than the simple cell (Figure 8), as reported previously (Rust et al., 2005). All subunits for a given cell had similar structure in space and time, but were translated relative to each other (Figure 8 A, B, top rows), as confirmed by the very similar frequency spectrum (Figure 8 A,B, bottom rows). For the complex cell, this is consistent with the hypothesis that it receives inputs from multiple simple cells at different locations (Hubel & Wiesel, 1962; Adelson & Bergen 1985, Vintch et al., 2015). Thus, the method presented here extends to V1 data, producing subunit estimates that are not constrained by assumptions of orthogonality (Rust et al., 2005) or convolutional structure (Vintch et al., 2015), but broadly resemble the results of previous work.

**Figure 8:**
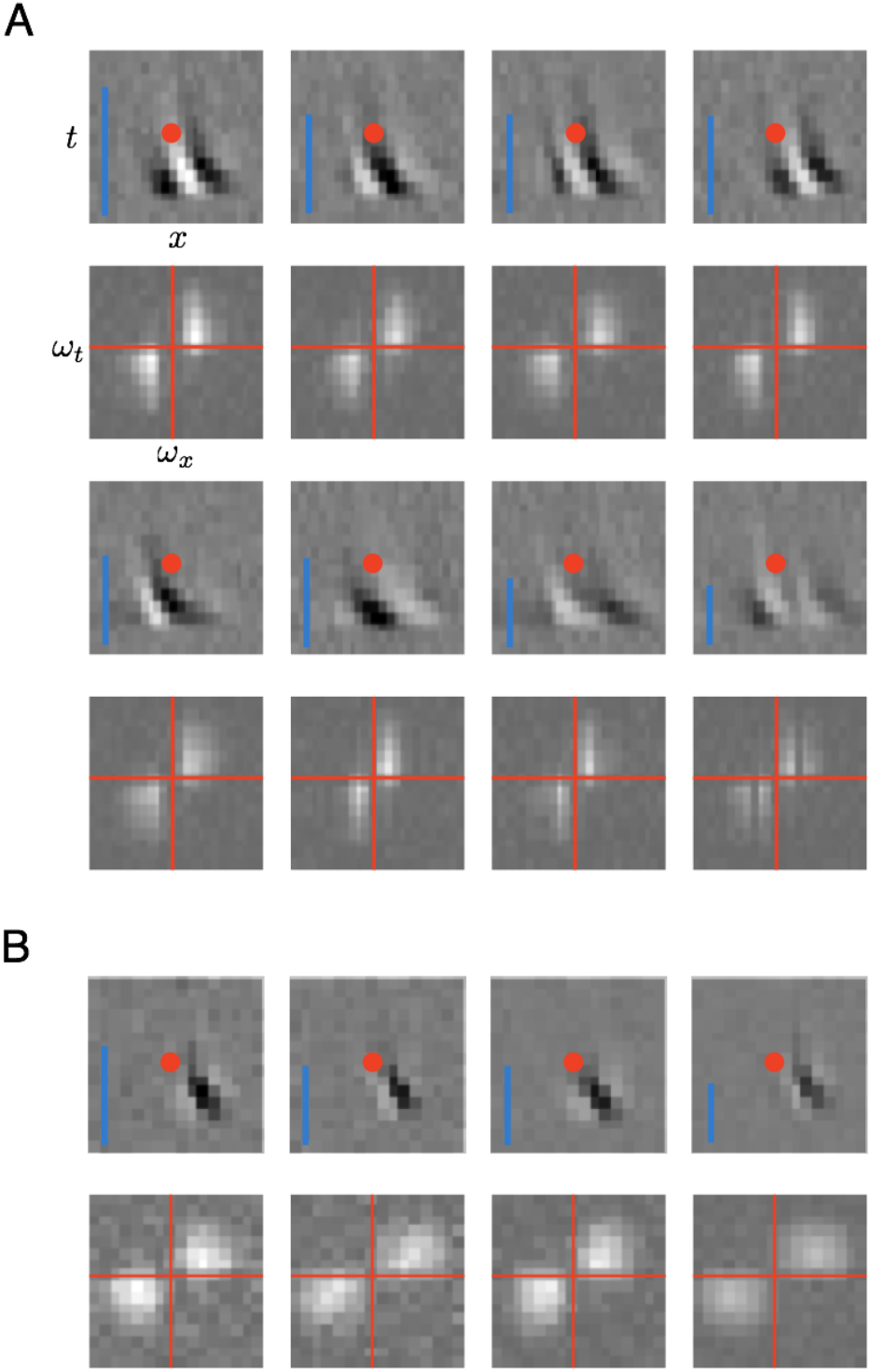
Application of subunit model to V1. (A) Estimated subunits using responses to flickering bar stimuli for the complex cell featured in Rust et al, 2005. Optimal number of subunits was found by cross-validation. Spatio-temporal filters (top row) and the fourier magnitude spectrum (bottom row) show translational invariance of estimated subunits. Red dots indicate center location for the spatio-temporal filters. Inset : Relative contribution to cell response for each subunit indicated by height of blue bar. (B) Same as (A) for the simple cell from Rust et al, 2005.

## Discussion

We developed an efficient modeling and estimation approach to study a ubiquitous motif of neural computation in sensory circuits: summation of responses of rectified subunits. The approach allowed efficient subunit estimation by clustering of spike-triggered stimuli, and the inclusion of priors for spatial localization to additionally constrain the fitting under conditions of limited data. The model and fitting procedure extended naturally to populations of cells, allowing joint fitting of shared subunits. We demonstrated the effectiveness of the method in capturing subunit computations in macaque parasol RGCs, specifically in stimulus conditions in which linear models completely fail, as well in V1 neurons. Below, we discuss the properties of the approach compared to others, and the implications of the application to retinal data.

### Modeling linear-nonlinear cascades

The subunit model is a natural generalization of linear-nonlinear (LN) cascade models, and the estimation procedure a natural generalization of the spike-triggered estimation procedures that enabled their widespread use in modeling neural computations. In the present model framework, a maximum-likelihood estimate of model parameters leads to a spike-triggered clustering algorithm for obtaining the early linear filters that represent model subunits. These linear components are the key, high-dimensional entities that define stimulus selectivity, and, in the absence of a robust computational framework, would be difficult to estimate.

Related prior work (Liu et al., 2017) developed a spike-triggered non-negative matrix factorization (SNMF) method for estimating subunits, and applied it to recordings from salamander RGCs. The estimated subunits were shown to be well-matched to independent physiological measurements of bipolar cell receptive fields, an impressive validation of the method. In comparison to the method presented here, there are several technical issues worth noting. First, the assumptions of the SNMF algorithm are not directly matched to the structure of the subunit response model: the present model assumes nonnegative subunit outputs, while the SNMF factorization method assumes that the subunit filters themselves are nonnegative. The assumption of nonnegative filters is inconsistent with suppressive surrounds previously reported in retinal bipolar cells (Dacey et al., 2000; Fahey & Burkhardt 2003; Turner et al., 2018), and limits the application of the method to other systems such as V1 (Figure 8). However, a recent modification of the SNMF method applies non-negativity constraints on subunit weights instead of the subunit filters (Jia et al., 2018), making it more suitable for other applications. Second, the SNMF method takes a sequence of additional steps to relate the estimated matrix factors to the parameters of an LNLN model. However in the present work, clustering of spike-triggered stimulus is shown to be equivalent to optimizing the prediction accuracy of a LNLN model, eliminating extra steps in relating model parameters to biology. Third, the comparison to bipolar cell receptive fields in the above studies relied on a regularization parameter that trades off sparseness and reconstruction error. This regularization can influence the size of the estimated subunits, and thus the comparison to bipolar receptive fields. In contrast, the present method includes a regularizer that encourages spatial contiguity of receptive fields, while exerting minimal influence on the size of estimated subunits, and is chosen using a cross-validation procedure.

Another study (Freeman et al., 2015) presented an approach to estimating subunits from OFF midget RGCs with visual inputs presented at single cone resolution, allowing a cellular resolution dissection of subunits. However, this approach assumed subunits that receive inputs from non-overlapping subsets of cones, which may not be an accurate assumption for some RGC types, including parasol cells. Moreover, the cone to subunit assignments were learned using a greedy procedure, which was effective for subunits composed of 1-2 cones, but is more likely to get stuck in local minima in more general conditions.

Recent studies fitted a large class of LNLN models to simulated RGC responses (McFarland et al., 2013), to recorded ON-OFF mouse RGC responses (Shi et al., 2018), and to flickering bar responses in salamander RGCs (Maheswaranathan et al., 2018) using standard gradient descent procedures. These approaches have the advantage of flexibility and simplicity. In contrast, we assume a specific subunit nonlinearity that leads to a specialized efficient optimization algorithm, and apply this to recorded data from primate RGCs. The choice of an exponential subunit non-linearity enables likelihood maximization for Gaussian stimuli, which leads to more accurate estimates with limited data, and reduces the computational cost of each iteration during fitting because the loss depends only on the collection of stimuli that elicit a spike (Ramirez et al., 2014). Additionally, the soft-clustering algorithm requires far fewer iterations than stochastic gradient descent algorithms used in fitting deep neural networks (Kingma et al, 2014; Duchi et al., 2011). Depending on the experimental conditions (e.g. type of stimuli, duration of recording, number of subunits to be estimated), the present approach may provide more accurate estimates of subunits, with less data, and more quickly.

Other recent studies applied convolutional and recurrent neural network models, fitted to natural scene responses, to mimic a striking variety of retinal response properties (McIntosh et al., 2016, Batty et al., 2016). The general structure of the model – convolutions and rectifying nonlinearities – resembles the subunit model used here. However, the number of layers and the interactions between channels do not have a direct correspondence to the known retinal architecture, and to date the specific properties of the inferred subunits, and their contributions to the behaviors of the model, have not been elucidated. The complexity of the underlying models suggest that a comparison to the biology could be difficult.

In the context of V1 neurons, the present approach has advantages and disadvantages compared to previous methods. Spike-triggered covariance (STC) methods (Rust et al., 2005) estimate an orthogonal basis for the stimulus subspace captured by the subunits. However, the underlying subunits may not be orthogonal, and even if they are, are not guaranteed to align with the basis elements. The method also requires substantial data to obtain accurate estimates of the basis. To reduce the data requirements, other methods (Vintch et al, 2015; Wu et al., 2015) imposed an assumption of convolutional structure on the subunits. In comparison, the method presented here does not assume orthogonal or convolutional subunits. Instead, approximate translational invariance emerges from data, consistent with known physiological properties of V1 neurons.

A generalization of the STC approach (Sharpee et al., 2004; Paninski 2003) involves finding a subspace of stimulus space which is most informative about a neuron’s response. This approach also yields a basis for the space spanned by subunits, but the particular axes of the space may or may not correspond to the subunits. Unlike the method presented here, this approach requires minimal assumptions about stimulus statistics, making it very flexible. However, methods based on information theoretic measures typically require much more data than are available in experimental recordings (Paninski 2003).

Although constraints on the structure of the subunit model (e.g. convolutional), and the estimation algorithm (e.g. spike-triggered covariance or clustering), both influence the efficiency of subunit estimation, the choice of priors used to estimate the subunits efficiently with limited data (i.e. regularization) is also important. In the present work, a regularizer was developed that imposes spatial contiguity, with minimal impact on the size of estimated subunits. A variety of other regularization schemes have been previously proposed for spatial contiguity for the estimation of V1 subunits (Park and Pillow, 2011). These could be incorporated in the estimation of the presented model using a corresponding projection operator.

As suggested by the results, another prior that could improve the subunit estimation is the hierarchical variant of the clustering approach (Figure S2). By splitting one of the subunits into two subunits at each step, models with different number of subunits can be estimated with greater efficiency, and preliminary exploration indicates that this may offer improvements in data efficiency and speed. However, potential tradeoffs emerging from the additional assumptions will require further study.

### Revealing the impact of non-linear computations using null stimuli

The null stimulus methodology revealed the failure of LN models, and isolated the nonlinear behaviors arising from receptive field subunits. Previous studies have developed specialized stimuli to examine different aspects of the cascaded linear-nonlinear model. Contrast reversing gratings (CRGs) have been the most commonly used stimuli for identifying the existence of a spatial nonlinearity. Schwartz & Rieke 2012 extend this logic to more naturalistic stimuli, with changes in responses to translations and rotations of texture stimuli revealing the degree of spatial nonlinearity. Bölinger et al., 2012 developed a closed-loop experimental method to measure subunit nonlinearities, by tracing the intensities of oppositely signed stimuli on two halves of the receptive field to measure iso-response curves. Freeman et al., 2015 developed stimuli in closed loop that targeted pairs of distinct cones in the receptive field, and used these to identify linear and nonlinear summation in RGCs depending on whether the cones provided input to the same or different subunits.

The null stimulus approach introduced here offers a generalization of the preceding methods. Null stimuli can be generated starting from any desired stimulus ensemble (e.g., natural images), and the method does not assume any particular receptive field structure (e.g. radial symmetry). Construction of the null stimuli does, however, require fitting a linear (or LN) model to the data in closed loop. Projecting a diverse set of stimuli (e.g. white noise) onto the null space yields a stimulus ensemble with rich spatio-temporal structure that generates diverse neural responses. By construction, the null stimulus is the closest stimulus to the original stimulus that is orthogonal to the receptive field, and thus minimally alters other statistical properties of the stimulus that affect neural response. This property is useful for studying the effect of spatial nonlinearities in the context of specific stimuli, such as natural scenes. Whereas contrast reversing gratings do not reveal stronger nonlinearities in OFF parasols compared to ON parasols in responses to natural stimuli (Turner and Rieke 2017), the null stimulus captures these differences (Figure 6), highlighting the advantages of tailored stimulation with rich spatio-temporal responses.

### Further applications and extensions

The assumptions behind the proposed subunit model and the application to primate RGC data enable efficient fitting and interpretability, but also lead to certain limitations that could potentially be overcome. First, the choice of exponential subunit nonlinearity improves efficiency by allowing the maximum expected likelihood approximation. Other nonlinearities that allow this approximation, such as rectified quadratic (Bölinger et al., 2012), could be explored. Second, the assumption of space-time separable RGC subunits significantly reduces the number of parameters that need to be estimated, but could be relaxed with more data. For example, a rank 2 approximation that allows for separate space-time separable center and surround filters for each subunit may be useful (Schweitzer-Tong, Enroth-Cugell & Pinto, 1970), given the center-surround structure of bipolar cell receptive fields (Dacey et al., 2000, Turner et al., 2018).

The methods developed here may also prove useful in tackling additional challenging problems in neural circuitry and nonlinear coding, in the retina and beyond. First, applying the model to high-resolution visual stimulation of the retina could be used to reveal how individual cones connect to bipolar cells, the likely cellular substrate of subunits, in multiple RGC types (see Freeman et al., 2015). Second, the incorporation of noise in the model, along with estimation of shared subunits (Figure 4), could help to probe the origin of noise correlations in the firing of nearby RGCs. Third, extending the subunit model to include gain control, another ubiquitous nonlinear computation in retina and throughout the visual system, could provide a more complete description of the responses in diverse stimulus conditions (Heeger, 1992; Carandini & Heeger 2011). In addition to scientific advances, a more accurate functional model is critical for the development of artificial retinas for treating blindness. Finally, the successful application of the method to V1 data (Figure 8) suggests that the method could be effective in capturing computations in other neural circuits that share the nonlinear subunit motif.

## Methods

### Recordings

Detailed preparation and recording methods are described elsewhere (Litke et al., 2004; Frechette et al., 2005; Chichilnisky and Kalmar, 2002). Briefly, eyes were enucleated from 7 terminally anesthetized macaque monkeys (Macaca sp.) used by other experimenters in accordance with institutional guidelines for the care and use of animals. Immediately after enucleation, the anterior portion of the eye and the vitreous were removed in room light. The eye was stored in darkness in oxygenated Ames’ solution (Sigma, St. Louis, MO) at 33°C, pH 7.4. Segments of isolated or RPE-attached peripheral retina (6-15mm temporal equivalent eccentricity (Chichilnisky and Kalmar, 2002), approximately 3mm x 3mm) were placed flat, RGC side down, on a planar array of extracellular microelectrodes. The array consisted of 512 electrodes in an isosceles triangular latticewith 60 μm inter-electrode spacing in each row and covering a rectangular region measuring 1800 μm x 900 μm. While recording, the retina was perfused with Ames’ solution (35°C for isolated recordings and 33°C for RPE-attached recordings), bubbled with 95% O2 and 5% CO_2_, pH 7.4. Voltage signals on each electrode were bandpass filtered, amplified, and digitized at 20 kHz.

A custom spike-sorting algorithm was used to identify spikes from different cells (Litke et al., 2004). Briefly, candidate spike events were detected using a threshold on each electrode, and voltage waveforms on the electrode and nearby electrodes in the 5ms period surrounding the time of the spike were extracted. Candidate neurons were identified by clustering the waveforms using a Gaussian mixture model. Candidate neurons were retained only if the assigned spikes exhibited a 1 ms refractory period and totaled more than 100 in 30 min of recording. Duplicate spike trains were identified by temporal cross-correlation and removed. Manual analysis was used to further select cells with a stable firing rate over the course of the experiment and with a spatially localized receptive field.

### Visual Stimuli

Visual stimuli were delivered using the optically reduced image of a CRT monitor refreshing at 120 Hz and focused on the photoreceptor outer segments. The optical path passed through the mostly transparent electrode array and the retina. The relative emission spectrum of each display primary was measured with a spectroradiometer (PR-701, PhotoResearch) after passing through the optical elements between the display and the retina. The total power of each display primary was measured with a calibrated photodiode (UDT Instruments). The mean photoisomerization rates for the L, M, and S cones were estimated by computing the inner product of the primary power spectra with the spectral sensitivity of each cone type, and multiplying by the effective collecting area of primate cones (0.6 μm2 (Angueyra and Rieke, 2013; Schnapf et al., 1990). During white noise and null stimulus, the mean background illumination level resulted in photoisomerization rates of (4100,3900,1600) for the (L,M,S) cones. The pixel size was either 41.6 microns (8 monitor pixels on a side) or 20.8 microns (4 monitor pixels on a side). A new white noise frame was drawn at refresh rates of 60 Hz (figure 2) or 30 Hz (Figure 5, 6). The pixel contrast (difference between the maximum and minimum intensities divided by the sum) was either 48% or 96% for each display primary, with mean contrast of 50%.

The details of natural stimuli are presented in Heitmann et al., 2016. Briefly, the stimuli consisted of images from the Van Hateren database, shown for one second each, jittered according to the eye movement trajectories recorded during fixation by awake macaque monkeys (Z.M. Hafed and R.J. Krauzlis, personal communication), (Heitmann et al, 2016, van Hateren and van der Schaaf, 1998). Image intensities produced mean photoisomerization rates of 1900, 1800, 700 for the L, M, S cones, respectively. The pixel width was 10.4 microns (2 monitor pixels on a side), and image frames refreshed at 120Hz. The training data consisting of 59 groups of 60 distinct natural scenes was interleaved with identical repeats of testing data consisting of 30 distinct natural scenes.

### Subunit Estimation

The light responses of RGCs were modeled using a cascade of two linear-nonlinear (LN) stages, followed by a Poisson spike generator. Specifically, the instantaneous stimulus-conditional firing rate was modeled as 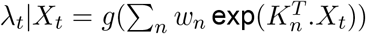 where *X_t_* is the visual stimulus at time *t, K_n_* are the spatial subunit filters, *w_n_* are non-negative weights on different subunits and 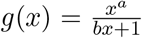, (*b* ≥ 0) is the output nonlinearity.

The parameters {*K_n_,w_n_,a,b*} are estimated by minimizing their negative log-likelihood given observed spiking responses *Y_t_* to visual stimuli *X_t_*, in two steps. In the first step, the parameters {*K_n_,w_n_*} are estimated using a clustering procedure while ignoring the nonlinearity (*g*). In the second step, {*g,w_n_*} are estimated using gradient descent, with the *K_n_* held fixed (up to a scale factor).

The negative log-likelihood for the first step can be simplified as follows:

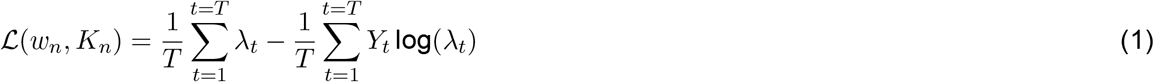

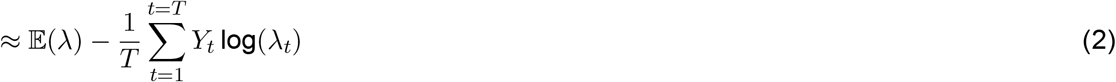

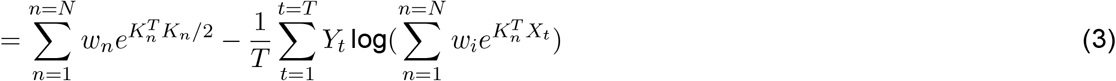

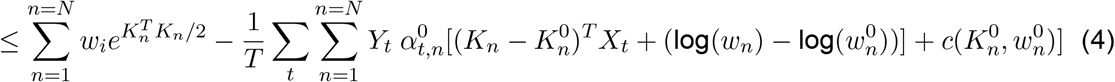

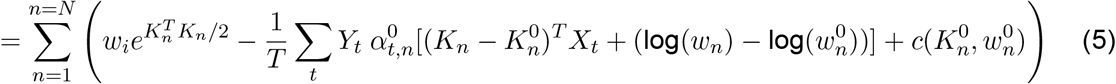

where 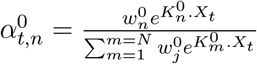. Since the first term in the log-likelihood (Eq. 1) does not depend on recorded responses (*Y_t_*), it is replaced with its expected value across stimuli (Eq. 2). Assuming the projections of the input distribution onto filters *K_n_* are approximately Gaussian, this expectation can be computed using the moment generating function of a Gaussian distribution (Eq. 3). The second term only depends on the stimuli for which the cell generated spikes (*Y_t_* > 0), reducing computational cost. Finally, to optimize the resulting non-convex function, a convex upper bound was minimized at each step. Specifically, the second term is replaced by a first-order Taylor approximation (Eq. 4) (the first term is already convex). Since the upper bound is tight at current parameter estimates, the successive convex optimization reduces the approximation of negative log-likelihood monotonically, leading to fewer iterations for convergence. This upper bound is separable across parameters of different subunits, as can be seen by re-arranging the summation over time and subunits (Eq. 5).

The minimization of the convex upper bound (Eq. 5) can accomplished using a clustering procedure composed of three sub-steps:

1. Update subunit activations (cluster weights) for each stimulus : 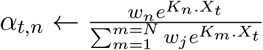.
2. Update linear subunit filters (cluster centers) : 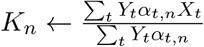.
3. Update relative weights for different subunits : 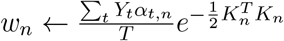.

The resulting subunits can be interpreted as a decomposition of the spike triggered average (STA), since the sum of estimated subunits (*K_n_*), weighted by their expected contribution to the response 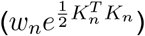, is proportional to the spike-triggered average:

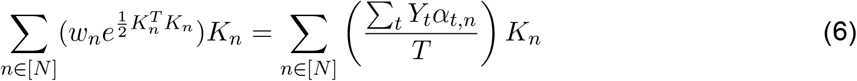

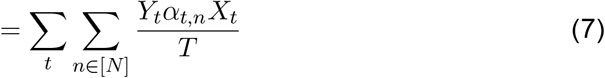

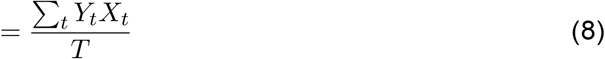

where (6) comes form (3) (at convergence), (7) comes from (2) (at convergence), and (8) arises because the *α_t,n_* sum to one. Step (6) may be interpreted as replacing the average spike rate, multiplied by contribution of each subunit. Step (7) may be interpreted as a weighted average of spike-triggered stimuli, where the weighting assigns responsibility for each spike to the subunits.

### Incorporating priors

The log-likelihood can be augmented with a regularization term, to incorporate prior knowledge about the structure of subunits, thereby improving estimation in the case of limited data. Regularization is incorporated into the algorithm by projecting the subunit kernels (*K_n_*) with a proximal operator (*P*_λ_), derived from the regularization function, after each update of the cluster centers (Figure 9). For a regularization term of the form λ*f*(*K*), the proximal operator is defined as 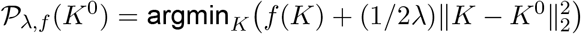. For each result (Figures 3, 4) the regularization strength (λ) was chosen through cross-validation (see below).

**Figure 9:**
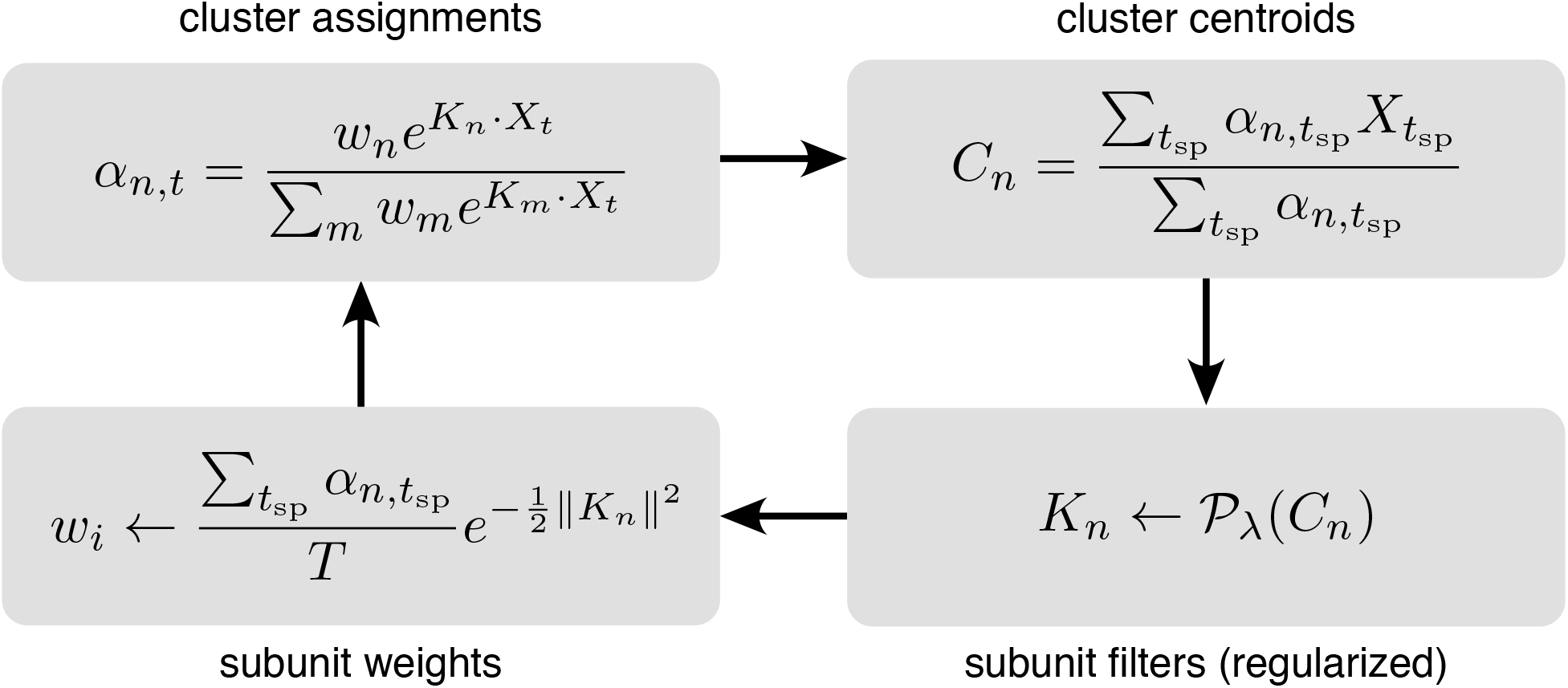
Iterative fitting of subunits, partitioned into four steps. The subunit kernels (*K_n_*) and weights (*w_n_*) are randomly initialized, and used to compute soft cluster assignments (*α_n,t_* upper left), followed by cluster centroid computation (*C_n_* - upper right), estimation of subunit kernels (*K_n_* - lower right), and subunit weights (*w_n_* - lower left). The summations are only over times when the cell generated a spike (*t*_sp_).

For *L*_1_-norm regularization (section 3), *f*(*K*) = Σ_*i*_|*K^i^*|, where *K^i^* is the value of *i*th pixel, and the proximal operator is a soft-thresholding operator 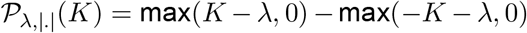. For the locally normalized *L*_1_-norm regularization, the proximal operator of the Taylor approximation was used at each step. The Taylor approximation is a weighted *L*_1_-norm with the weight for each pixel determined by the parameter value of the neighboring pixels. Hence, the proximal operator for *i*th pixel is 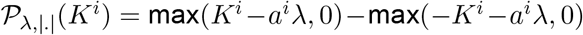, where *a^i^* = 1/(*ϵ* + Σ_*j*∈Neighbors(*i*)_ |*K^j^*|) for small *ϵ* = 0.01. Simulations were used to verify that this regularizer induces a preference for spatially contiguous subunits, while being relatively insensitive to subunit area (compared with *L*_1_ regularization).

The entire procedure is summarized in Figure 9, with simplified notation illustrating the four key steps.

### Subunit estimation using hierarchical clustering

As shown in Figure 2, a hierarchical organization of subunits was observed in fits to RGC data, with one subunit broken down into two subunits each time the total number of subunits was increased by one. This hierarchical structure can be enforced explicitly to make the estimation procedure more efficient.

Since the softmax weights *α_m_* (between 0 and 1) can be interpreted as the ‘activation probability’ of subunit m, hierarchical clustering is performed by estimating two children subunits *m*_1_,*m*_2_ with factorized activation probability : *α*_*m*1_ = *α_m_α*_*m*1|*m*_, where *α*_*m*1|*m*_ is the conditional probability of child *m*_1_ given the activation of parent subunit with 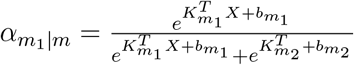. The factorization is accurate if the activation of the parent subunit is the sum of activation of the children subunits (*e^K_m_.X + b_m_^* = *e*^*K*_*m*1_.*X* + *b*_*m*1_^ + *e*^*K*_*m*_2__.*X*+*b*_*m*_2__^).

The children subunits were initialized by adding independent random noise to parent subunits (*K_m_i__* = *K_m_* + *ϵ_i_*) and equally dividing the subunit weight into two 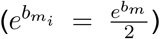. Hierarchical clustering is performed by iteratively updating the cluster assignments (*α_m_i__*) and cluster centers *(K_mi_, b_mi_*) for the children subunits using the factored *α* as outlined above.

The parent subunit for splitting is chosen greedily, to give maximum improvement in log-likelihood at each step. Hierarchical subunit estimation provides a computationally efficient way to estimate the entire solution path with different *N*. The estimated subunits obtained with this procedure are shown in Figure S2.

### Population model

In Section 4, a method to estimate a common bank of subunits jointly using multiple nearby cells is described. For each cell *c*, the firing rate is given by 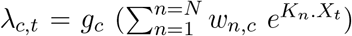 where *w_n,c_* are the cell specific subunit weights. The model parameters are estimated by maximizing the summation of log-likelihood across cells.

Similar to the single cell case, the estimation is performed in two steps: 1) Ignoring the output nonlinearity *g_c_*, estimate {*K_n_,w_n,c_*} by clustering, and 2) Estimate *g_c_, w_n,c_* and the magnitude of *K_n_* for each cell independently, using gradient descent with vector direction of {*K_n_*} fixed up to a scalar. Clustering in the first step can be interpreted as finding a common set of centroids to cluster the spike triggered stimuli for multiple cells simultaneously. The following steps are repeated iteratively:

1. Update subunit activations (cluster weights) for each stimulus and cell :

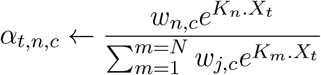
2. Update linear subunit filters (cluster centers) using population responses :

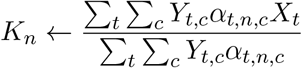
3. Update subunit weights (*w_n,c_*) for each cell :

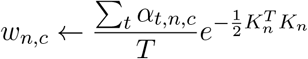

### Application to neural responses

The subunit estimation algorithm was applied to a spatial segment of stimulus around the receptive field (5.2 micron boundary) and to the RGC spike counts over time (8.33ms bin size). The data were partitioned into three groups: testing data consisted of contiguous segment of data (last 10%) and rest of the data was randomly partitioned for training (90%) and validation (10%). Models were fit with different numbers of subunits *N* (1-12) and regularization values λ. Hyperparameter selection (*N* and λ) was performed by averaging the performance of multiple fits with different training/validation partitions and random initializations (see Figure captions for details). For a given number of subunits, the regularization value was chosen by optimizing the performance on validation data. Similarly, the most accurate model was chosen by optimizing over the number of subunits as well as regularization strength. When data with repeated presentations of the same stimulus were available (e.g. Figure 6, 7), the prediction accuracy was evaluated by measuring the correlation between peri-stimulus time histogram (PSTH) of recorded data and the predicted spikes for the same number of repetitions. The PSTH was computed by averaging the discretized responses over repeats and gaussian smoothing with standard deviation 16.66ms.

For null stimulus, responses from 30 repeats of a 10 sec long null stimulus were divided into training (first 5 sec) and testing (last 5 sec). Since the training data were limited, the subunit filters and weights (phase 1) were first estimated from long white noise stimulus and only the subunit scales and output nonlinearity (phase 2) were estimated using the null stimulus.

For natural scenes stimuli, the spatial kernel for subunits was first estimated from responses to the white noise stimulus (phase 1), and the scales of subunits and output nonlinearity were then fit using responses to a natural stimulus (phase 2). This procedure was used because subunits estimated directly from natural scenes did not show any improvement in prediction accuracy, and all the subunits were essentially identical to the RF, presumably because of the strong spatial correlations in natural images.

Previously published V1 responses to flickering bars were used for fitting subunits (see Rust et al., 2015 for details). In contrast to the retinal data, the two dimensions of the stimulus were time and one dimension of space (orthogonal to the preferred orientation of the cell). The number of subunits was chosen by cross validation as described above.

### Simulated RGC model

To relate estimated subunits to bipolar cells, the model was fitted to responses generated from a simulated RGC (Appendix I). In the simulation, each RGC received exponentiated inputs from bipolar cells, which in turn linearly summed inputs from cones. The spatial dimensions of the simulation and the number of cones and bipolar cells were approximately matched to parasol RF center at the eccentricities of the recorded data (Schwartz and Rieke, 2011; Jacoby et al., 2000). Below, all the measurements are presented in grid units (g.u.), with 1 g.u. = 1 micron. The first layer consisted of 64 cones arranged on a jittered hexagonal lattice with nearest-neighbor distance 180 g.u., oriented at 60 degrees with respect to the stimulus grid. The location of each cone was independently jittered with a radially symmetric Gaussian with standard deviation 12.6 g.u. Stimuli were pooled by individual cones with a spatio-temporally separable filter, with time course derived from a typical parasol cell and a Gaussian spatial filter (standard deviation = 12 g.u.). The photoreceptor weights were chosen to give a mean firing rate of ≈ 19 spikes/sec.

The second layer of cascade consisted of 12 bipolar cells, each summing the inputs from a spatially localized set of photoreceptors, followed by an exponential nonlinearity. The photoreceptors were assigned to individual bipolars by k-means clustering (12 clusters) of their locations. Finally, the RGC firing rate is computed by a positively weighted summation of bipolar activations. The bipolar weights were varied within 10% of the mean weight.

The subunit model was fitted on 1.5 hour long white noise stimulus such that the receptive field was roughly covered by 6 x 6 stimulus pixels, as in the recorded experimental data (Figure 2).

### Null stimulus

We constructed null stimuli to experimentally validate the estimated subunits (Figures 5, 6). For a spatio-temporal stimulus movie, the goal is to find the closest stimulus such that the convolution with a spatio-temporal linear filter gives 0. For simplicity, we describe the algorithm only for a single cell, and the extension to multiple cells is straightforward. Let *a*(*x, y, t*) represent the spatiotemporal linear filter, estimated by computing STA on white noise responses. The orthogonality constraint for a null stimulus *S*(*x, y, t*) is written as:

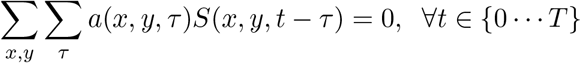

.The constraints for successive frames are not independent, but they become so when transformed to the temporal frequency domain. Writing *a*(.,., *ω*) and *S*(.,., *ω*) for the Fourier transform of *a*(.,.,*t*) and *S*(.,.,*t*) respectively, the constraints may be rewritten as:

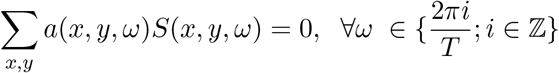

The spatial content can be projected onto the space orthogonal to the estimated spatial receptive field, for each temporal frequency. Specifically, given a vectorized stimulus frame *S_w_* and a matrix *A* with columns corresponding to vectorized receptive fields of a set of cells, the null frame is obtained by solving the following optimization problem:

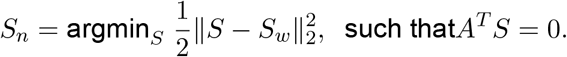

The solution may be written in closed form:

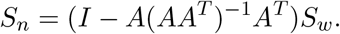

The receptive field for each cell was estimated by computing a space-time separable approximation of the spike-triggered average (STA), with the support limited to the pixels with value significantly different from background noise (absolute value of pixel value > 2.5 *σ*).

In addition to the orthogonality constraint, two additional constraints are imposed: a) each pixel must be in the range of monitor intensities (0-225) and b) the variance of each pixel over time must be preserved, to avoid contrast adaption in photoreceptors. These two constrains were incorporated using Dykstra’s algorithm (Boyle and Dykstra, 1986), which iteratively projects the estimated null stimulus into the subspace corresponding to individual constraints until convergence. Setting the contrast of initial white noise to 50% was necessary to satisfy these additional constraints. Finally, the pixel values of the resulting null stimuli were discretized to 8-bit integers, for display on a monitor with limited dynamic range.

For space-time separable (rank-1) STA (as in the recorded RGC data), spatio-temporal nulling gives the same solution as spatial-only nulling. To see this, assume *a*(*x,y,t*) = *a*_1_(*x,y*)*a*_2_(*t*), a space-time separable linear filter. Hence, the null constraint can be re-written as

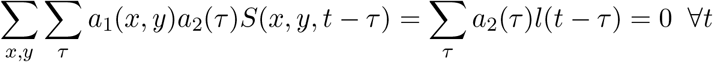

where *l*(*t* − *t*) = Σ_*x,y*_ *a*_1_(*x, y*)*S*(*x, y,t* − *τ*). In the frequency domain this is written as

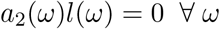

Since *a*_2_(*t*) has limited support in time, it has infinite support in frequency. Hence, *l*(*ω*) = 0 ∀*ω* implying spatial-only nulling *l*(*t*) = 0 ∀*t*.

## Acknowledgements

This work was supported by NSF IGERT Grant 0801700 (NB), NIH NEI F31EY027166 (CR), NSF GRFP DGE-114747 (CR, NB), Wu Tsai Neurosciences Institute Interdisciplinary Scholar Award (GG), Pew Charitable Trust Scholarship in the Biomedical Sciences (AS), Howard Hughes Medical Institute (ES), NIH NEI R01-EY021271 (EJC), NIH NEI P30-EY019005 (EJC). We thank Liam Paninski for helpful suggestions on the manuscript, Jill Desnoyer and Ryan Samarakoon for technical assistance, and H. Fox, M. Taffe, T. Albright, R. Krausliz, R. Siegel, K. Bankiewicz, C. Darian-Smith, J. Carmena, J. Wallis, E. Callaway, T. Moore, S. Morairty and the UC Davis Primate Center for providing access to retinas.

## Author contributions

NS, NB, CR, AK, GG collected the MEA data, NS, EJC, ES conceived and designed the experiments, NS implemented the subunit model and analyzed data, AS and AL developed and supported the MEA hardware and software, NS, EJC and ES wrote the paper, all authors edited the paper, EJC and ES supervised the project.

## Supplementary

### Validation of subunit estimation of simulated RGC responses

**Figure S1:**
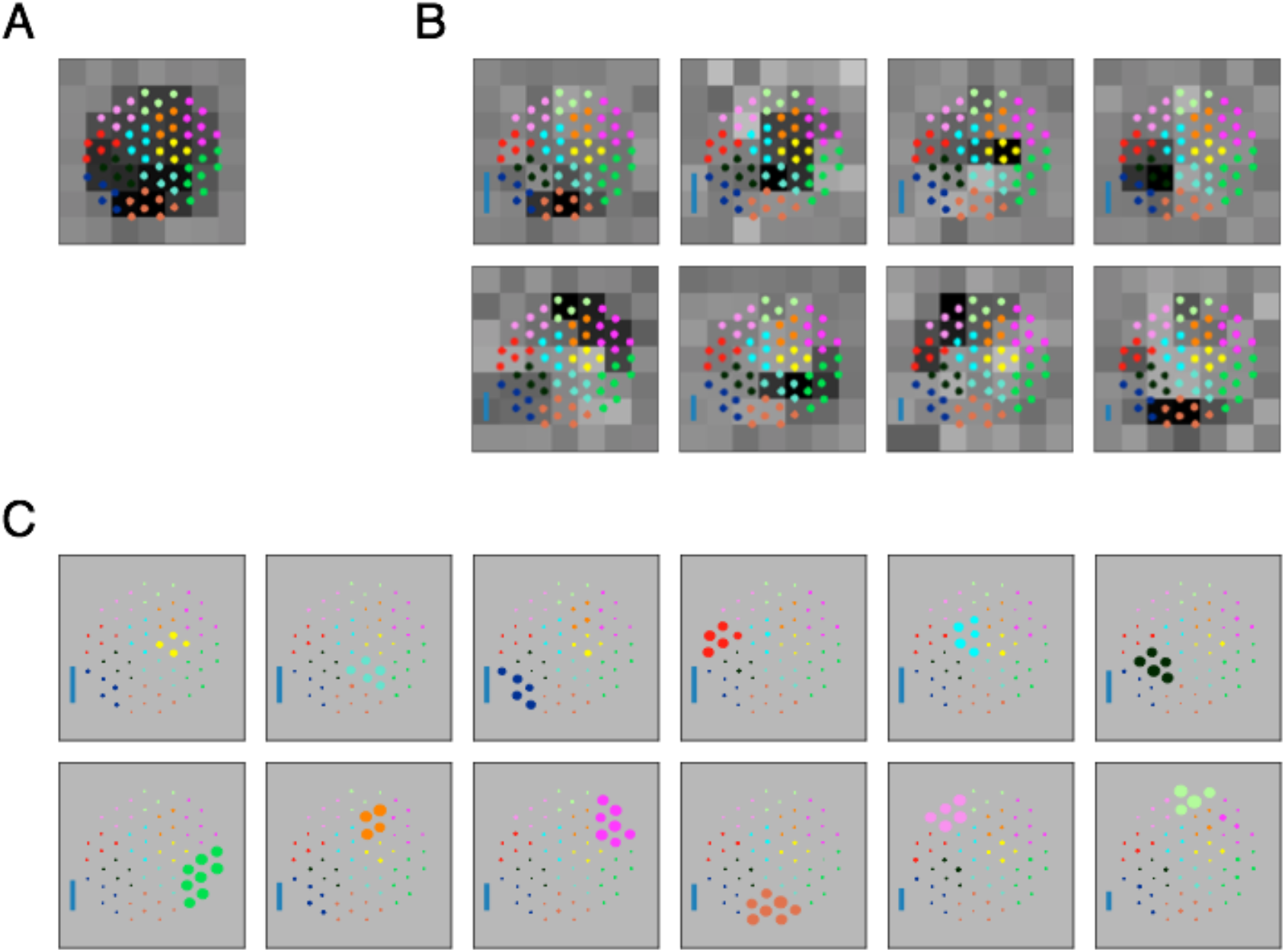
Validation of the subunit fitting algorithm on simulated RGC data. (A) Receptive field (spike-triggered average) of the simulated RGC with cascaded linear nonlinear units. Stimulus is temporally filtered by 64 photoreceptors, organized on a jittered hexagonal grid. Photoreceptor activations are summed by 12 bipolar cells, each connecting to 4-8 photoreceptors. Bipolar activations are exponentiated and added to give the poisson firing rate of the cell. Dots indicate photoreceptor locations, photoreceptors with same color connect to a single bipolar cell. See Methods for further details. (B) Subunits estimated from simulated RGC responses to 24 min of coarse white noise reveals fewer (8) subunits compared to the actual number of bipolar cells (12). The spatially localized subunits are aggregates of multiple, nearby underlying bipolar cells. The subunits are partially overlapping and tile to cover the receptive field. Inset: Blue bar height indicates the relative strength (average contribution to response over stimulus ensemble) of subunit. (C) Subunits estimated from simulated RGC responses to 6 hours of fine resolution white noise stimulus recovers the underlying bipolar cells. Size of dots indicate relative weights of different photoreceptors to the estimated subunits (12 total). Each subunit has strong inputs from photoreceptors connected to same bipolar cell. Inset: Same as B.

### Estimating subunits by hierarchical clustering

**Figure S2:**
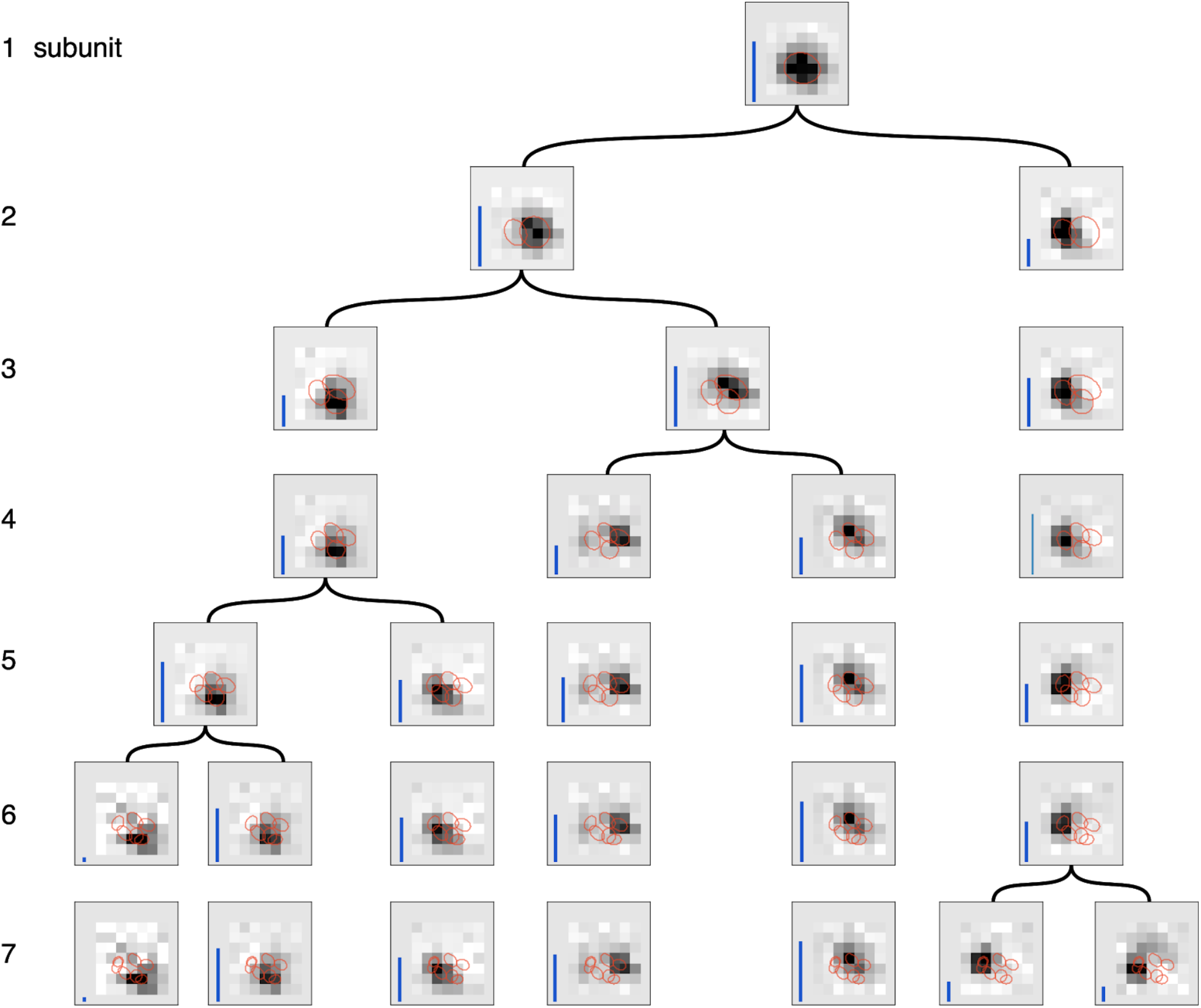
Gradual partitioning of the receptive field into subunits by hierarchical clustering. Different number of subunits (rows) estimated by splitting one parent subunit into two subunits at each step. Children subunits estimated by soft-clustering the simulated spikes of the parent subunit, with the simulated spikes for parent subunit computed as the spiking activity of the cell, weighed by its soft-max subunit activation (see Methods). The parent subunit that gives maximum decrease in log-likelihood on training data is chosen for splitting. The choice of training data, preprocessing and figure details same as Figure 2. The achieved splitting of subunits is similar to the pattern of splitting in Figure 2 for small number of subunits (1-5 subunits), but differs for larger number of subunits (6, 7 subunits). This suggests that enforcing the hierarchical constraint could lead to a more efficient estimation procedure.

